# Low-dimensional learned feature spaces quantify individual and group differences in vocal repertoires

**DOI:** 10.1101/811661

**Authors:** Jack Goffinet, Samuel Brudner, Richard Mooney, John Pearson

## Abstract

Increases in the scale and complexity of behavioral data pose an increasing challenge for data analysis. A common strategy involves replacing entire behaviors with small numbers of handpicked, domain-specific features, but this approach suffers from several crucial limitations. For example, handpicked features may miss important dimensions of variability, and correlations among them complicate statistical testing. Here, by contrast, we apply the variational autoencoder (VAE), an unsupervised learning method, to learn features directly from data and quantify the vocal behavior of two model species: the laboratory mouse and the zebra finch. The VAE converges on a parsimonious representation that outperforms handpicked features on a variety of common analysis tasks, enables the measurement of moment-by-moment vocal variability on the timescale of tens of milliseconds in the zebra finch, provides strong evidence that mouse ultrasonic vocalizations do not cluster as is commonly believed, and captures the similarity of tutor and pupil birdsong with qualitatively higher fidelity than previous approaches. In all, we demonstrate the utility of modern unsupervised learning approaches to the quantification of complex and high-dimensional vocal behavior.

## 1 Introduction

Quantifying the behavior of organisms is of central importance to a wide range fields including ethology, linguistics, and neuroscience. Yet given the variety and complex temporal structure of many behaviors, finding concise yet informative descriptions has remained a challenge. Vocal behavior provides a paradigmatic example: audio data are notoriously high dimensional and complex, and despite intense interest from a number of fields, and significant progress, many aspects of vocal behavior remain poorly understood. A major goal of these various lines of inquiry has been to develop methods for the quantitative analysis of vocal behavior, and these efforts have resulted in several powerful approaches that enable the automatic or semi-automatic analysis of vocalizations [48, 7, 50, 42, 49, 30, 32, 26, 18].

Key to this approach has been the existence of software packages that calculate acoustic features for each unit of vocalization, typically a syllable [4, 48, 50, 7, 6]. For example, Sound Analysis Pro, focused on birdsong, calculates 14 features for each syllable, including duration, spectral entropy, and goodness of pitch, and uses the set of resulting metrics as a basis for subsequent clustering and analysis [48]. More recently, MUPET and DeepSqueak have applied a similar approach to mouse vocalizations, with a focus on syllable clustering [50, 7]. Collectively, these and similar software packages have helped facilitate numerous discoveries, including circadian patterns of song development in juvenile birds [10], cultural evolution among isolate zebra finches [12], and differences in ultrasonic vocalizations (USVs) between mouse strains [50].

Despite these insights, this general approach suffers from several limitations: First, handpicked acoustic features are often highly correlated, and these correlations can result in redundant characterizations of vocalization. Second, an experimenter-driven approach may exclude features that are relevant for communicative function or, conversely, may emphasize features that are not salient or capture negligible variation in the data. Third, there is no diagnostic approach to determine when enough acoustic features have been collected: Could there be important variation in the vocalizations that the chosen features simply fail to capture? Lastly and most generally, committing to a syllable-level analysis necessitates a consistent definition of syllable boundaries, which is often difficult in practice. It limits the types of structure one can find in the data, and is often difficult to relate to time series such as neural data, for which the relevant timescales are believed to be orders of magnitude faster than syllable rate.

Here, we address these shortcomings by applying a data-driven approach based on variational autoencoders (VAEs) [25, 39] to the task of quantifying vocal behavior in two model species: the laboratory mouse (*Mus musculus*) and the zebra finch (*Taeniopygia guttata*). The VAE is an unsupervised modeling approach that learns from data a pair of probabilistic maps, an “encoder” and a “decoder,” capable of compressing the data into a small number of latent variables while attempting to preserve as much information as possible. In doing so, it discovers features that best capture variability in the data, offering a nonlinear generalization of methods like PCA and ICA that adapts well to high-dimensional data like natural images [8, 17]. By applying this technique to collections of single syllables, encoded as time-frequency spectrograms, we looked for latent spaces underlying vocal repertoires across individuals, strains, and species, asking whether these data-dependent features might reveal aspects of vocal behavior overlooked by traditional acoustic metrics and provide more principled means for assessing differences among these groups.

Our contributions are fourfold: First, we show that the VAE’s learned acoustic features outperform common sets of handpicked features in a variety of tasks, including capturing acoustic similarity, representing a well-studied effect of social context on zebra finch song, and comparing the USVs of different mouse strains. Second, using learned latent features, we report new results concerning both mice and zebra finches, including the finding that mouse USV syllables do not appear to cluster into distinct subtypes, as is commonly assumed, but rather form a broad continuum. Third, we present a novel approach to characterizing stereotyped vocal behavior that does not rely on syllable boundaries, one which we find is capable of quantifying subtle changes in behavioral variability on tens-of-milliseconds timescales. Lastly, we demonstrate that the VAE’s learned acoustic features accurately reflect the relationship between songbird tutors and pupils. In all, we show that data-derived acoustic features confirm and extend findings gained by existing approaches to vocal analysis, and offer distinct advantages over handpicked acoustic features in several critical applications.

## 2 Results

### 2.1 Variational autoencoders learn a low-dimensional space of vocal features

We trained a variational autoencoder (VAE) [25, 39] to learn a probabilistic mapping between vocalizations and a latent feature space. Specifically, we mapped single-syllable spectrogram images (*D* = 16, 384 pixels) to vectors of latent features (*D* = 32) and back to the spectrogram space (Figure 1a). As with most VAE methods, we parameterized both the encoder and decoder using convolutional neural networks, which provide useful inductive biases for representing regularly sampled data such as images or spectrograms. The two maps are jointly trained to maximize a lower bound on the probability of the data given the model (see Methods). As in other latent variable models, we assume each observed spectrogram can be explained by an unobserved “latent” variable situated in some “latent space.” As visualized in Figure 1b, the result is a continuous latent space that captures the complex geometry of vocalizations. Each point in this latent space represents a single spectrogram image, and trajectories in this latent space represent sequences of spectrograms that smoothly interpolate between start and end syllables (Figure 1c). Although we cannot visualize the full 32-dimensional latent space, methods like PCA and the UMAP algorithm [31] allow us to communicate results in an informative and unsupervised way. The VAE training procedure can thus be seen as a compression algorithm that represents each spectrogram as a collection of 32 numbers describing data-derived vocal features. In what follows, we will show that these features outperform traditional handpicked features on a wide variety of analysis tasks.

**Figure 1:**
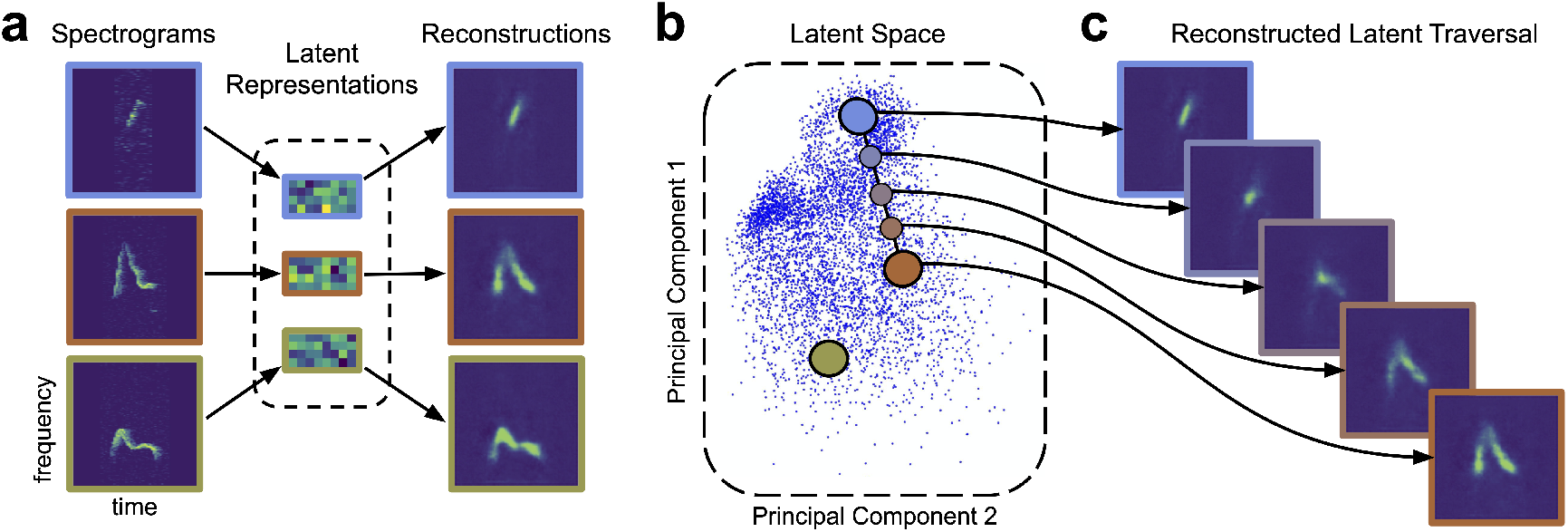
Variational autoencoders learn a latent acoustic feature space. **a)** The VAE takes spectrograms as input (left column), maps them via a probabilistic “encoder” to a vector of latent dimensions (middle column), and reconstructs a spectrogram via a “decoder” (right column). The VAE attempts to ensure that these probabilistic maps match the original and reconstructed spectrograms as closely as possible. **b)** The resulting latent vectors can then be visualized via dimensionality reduction techniques like principal components analysis. **c)** Interpolations in latent space correspond to smooth syllable changes in spectrogram space. A series of points (dots) along a straight line in the inferred latent space is mapped, via the decoder, to a series of smoothly changing spectrograms (right). This correspondence between inferred features and realistic dimensions of variation is often observed when VAEs are applied to data like natural images [25, 39]

Finally, we note that, while the VAE is compressive — that is, it discards some data — this is both necessary in practice and often desirable. First, necessity: as noted above, nearly all current methods reduce raw audio waveforms to a manageable number of features for purposes of analysis. This is driven in part by the desire to distill these complex sounds into a small collection of interpretable features, but it also stems from the needs of statistical testing, which suffers drastic loss of power for high-dimensional data without advanced methods. Second, this compression, as we will show, is often beneficial, as it facilitates visualization and analyses of large collections of vocalizations in new ways. Thus, we view the VAE and its compression-based approach as complementary to both other dimension-reduction techniques like PCA and traditional acoustic signal processing as a method for learning structure from complex data.

### 2.2 Learned features capture and expand upon typical acoustic features

Most previous approaches to analyzing vocalizations have focused on tabulating a predetermined set of features such as syllable duration or entropy variance that are used for subsequent processing and analysis [50, 7, 48, 4]. We thus asked whether the VAE learned feature space simply recapitulated these known features or also captured new types of information missed by traditional acoustic metrics. To address the first question, we trained a VAE on a publicly available collection of mouse USVs (31,440 total syllables [1]), inferred latent features for each syllable, and colored the results according to three acoustic features — frequency bandwidth, maximum frequency, and duration — calculated by the analysis program MUPET [50]. As Figures 2a-c show, each acoustic feature appears to be encoded in a smooth gradient across the learned latent space, indicating that information about each has been preserved. In fact, when we quantified this pattern by asking how much variance in a wide variety of commonly used acoustic metrics could be accounted for by latent features (see Methods), we found that values ranged from 64% to 95%, indicating that most or nearly all traditional features were captured by the latent space (see Figure S1 for individual acoustic features). Furthermore, we found that, when the analysis was reversed, commonly used acoustic features were not able to explain as much variance in the VAE latent features, indicating a prediction asymmetry between the two sets (Figure 2e). That is, the learned features carry most of the information available in traditional features, as well as unique information missed by those metrics.

**Figure 2:**
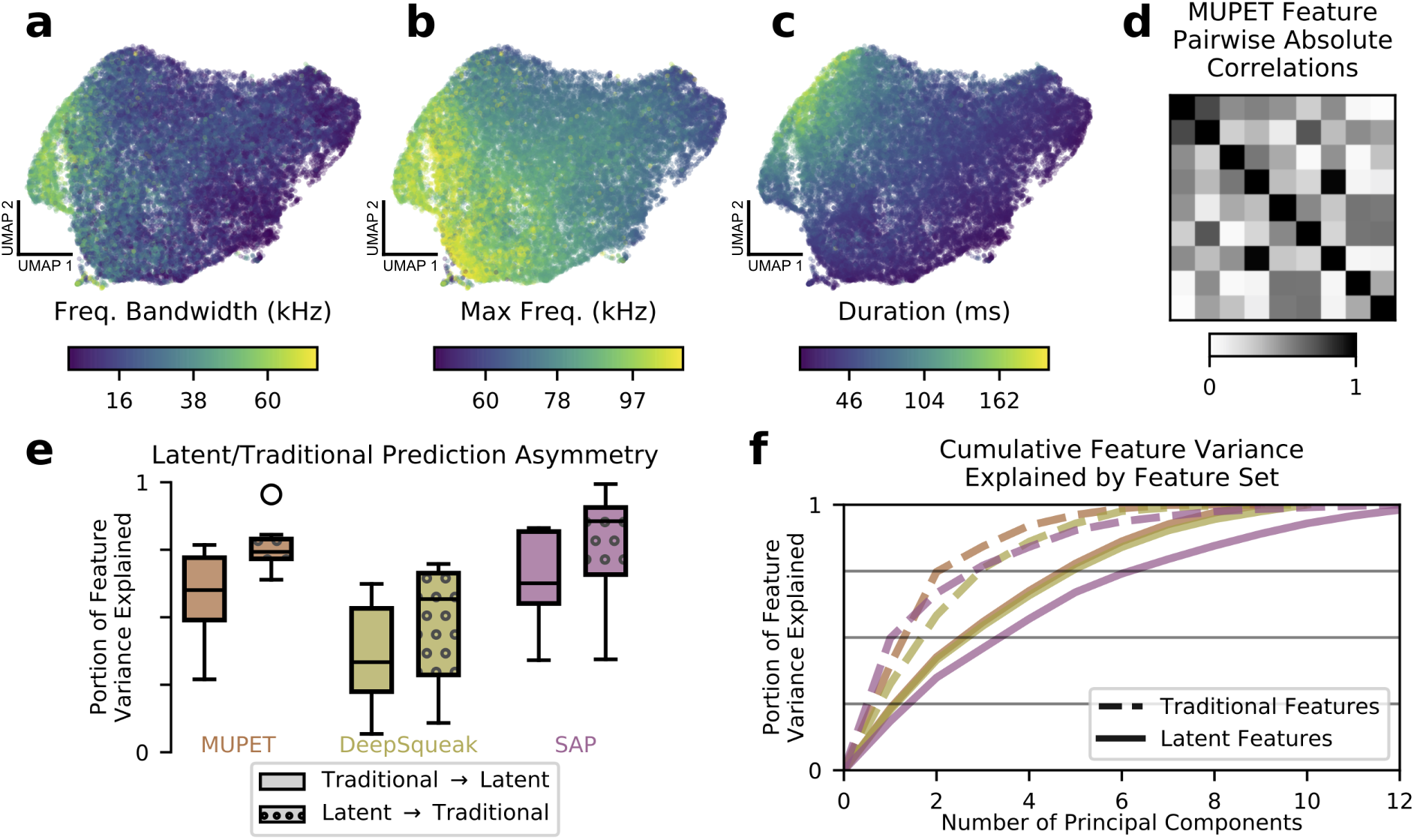
Learned acoustic features capture and expand upon traditional features. **a-c**) UMAP projections of latent descriptions of mouse USVs, colored by three traditional acoustic features. The smoothly varying colors show that these traditional acoustic features are represented by gradients within the latent feature space. **d)** Many traditional features are highly correlated. When applied to the mouse USVs from **a-c**, the acoustic features compiled by the analysis program MUPET have high correlations, effectively reducing the number of independent measurements made. **e)** To better understand the representational capacity of traditional and latent acoustic features, we used each set of features to predict the other and vice versa (see Methods). We find that, across software programs, the learned latent features were better able to predict the values of traditional features than vice-versa, suggesting they have a higher representational capacity. Central line indicates median, upper and lower box the 25^th^ and 75^th^ percentiles, respectively. Whiskers indicate 1.5 times the interquartile range. Feature vector dimensions: MUPET, 9; Deepsqueak, 10; SAP, 13; mouse latent, 7; zebra finch latent, 5. **f)** As a another test of representational capacity, we performed PCA on the feature vectors to determine the effective dimensionality of the space spanned by each set of features (see Methods). We find in all cases that latent features require more principal components to account for the same portion of feature variance, evidence that latent features span a higher dimensional space than traditional features applied to the same datasets. Colors are as in **e**. Latent features with colors labeled “MUPET” and “DeepSqueak” refer to the *same* set of latent features, truncated at different dimensions corresponding to the number of acoustic features measured by MUPET and DeepSqueak, respectively

We thus attempted to compare the effective representational capacity of the VAE to current best approaches in terms of the dimensionalities of their respective feature spaces. We begin by noting that the VAE, although trained with a latent space of 32 dimensions, converges on a parsimonious representation that makes use of only 5 to 7 dimensions, with variance apportioned roughly equally between these (Figure S3, [8]). For the handpicked features, we normalized each feature independently by z-score to account for scale differences. For comparison purposes, we applied the same normalization step to the learned features, truncated the latent dimension to the number of handpicked features, and calculated the cumulative feature variance as a function of number of principal components (Figure 2f). In such a plot, shallow linear curves are preferred, since this indicates that variance is apportioned roughly equally among principal components and the effective dimensionality of the space is large. Equivalently, this means that the eigenvalue spectrum of the feature correlation matrix is close to the identity. As Figure 2f thus makes clear, the spaces spanned by the learned latent features have comparatively higher effective dimension than the spaces spanned by traditional features, suggesting that the learned features have a higher representational capacity. While the three software packages we tested (SAP [48], MUPET [50], DeepSqueak [7]) measure upwards of 14 acoustic features per syllable, we find that these features often exhibit high correlations (Figure 2d,S2), effectively reducing the number of independent measurements made. While correlations among features are not necessarily undesirable, they can complicate subsequent statistical testing because nonlinear relationships among features violate the assumptions of many statistical tests. VAE features, which allow for nonlinear warping, avoid this potential difficulty.

The degree to which the learned features capture novel information can also be demonstrated by considering their ability to encode a notion of spectrogram similarity, since this is a typical use to which they are put in clustering algorithms (although see [50] for an alternative approach to clustering). We tested this by selecting query spectrograms and asking for the closest spectrograms as represented in both the DeepSqueak acoustic feature space and the VAE’s learned latent space. As Figure 3 shows, DeepSqueak feature space often fails to return similar spectrograms, whereas the learned latent space reliably produces close matches (see Figure S6 for comparisons to metrics in spectrogram space, Figure S7 for a representative sample using all feature sets, and see Figure S8 for more details on nearest neighbors returned by DeepSqueak feature space). This suggest that the learned features better characterize local variation in the data by more accurately arranging nearest neighbors.

**Figure 3:**
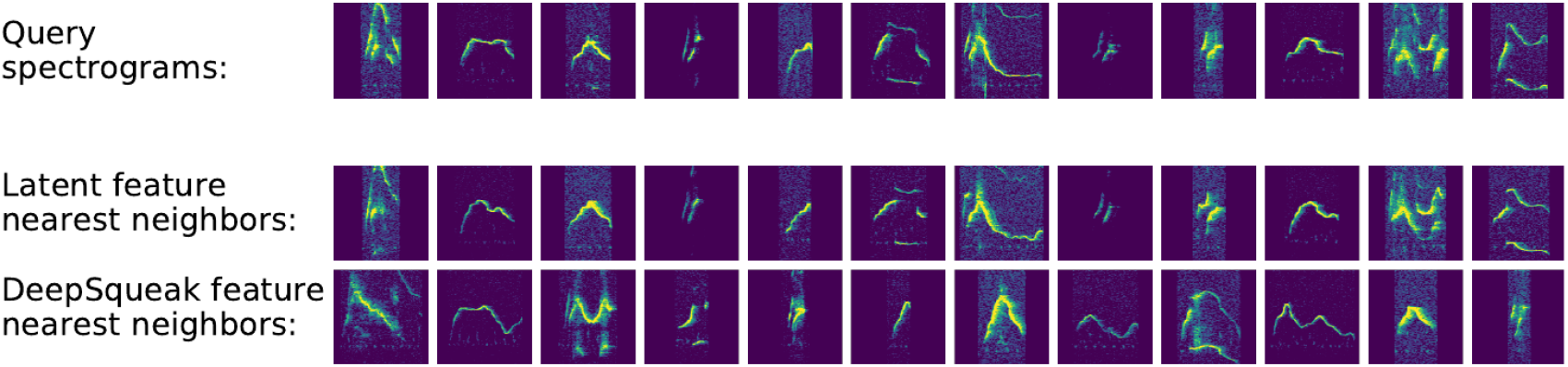
Latent features better represent acoustic similarity. **top row**: example spectrograms **middle row**: nearest neighbors in latent space **bottom row**: nearest neighbors in DeepSqueak feature space.

### 2.3 Latent spaces facilitate comparisons between vocal repertoires

Many experimental designs require quantifying differences between sets of vocalizations. As a result, the ability of a feature set to distinguish between syllables, individuals, and groups poses a key test of the VAE-based approach. Here, we apply the VAE latent features to several comparison problems for which handpicked features are often used.

A common comparison in birdsong research is that between female-directed and undirected song. It is well-established that directed song is more stereotyped and slightly faster than undirected song [46]. We thus asked whether the learned features could detect this effect. In Figure 4a we plot the first two principal components of acoustic features calculated by the Sound Analysis Pro software package [48] for both directed and undirected renditions of a single zebra finch song syllable. We note a generally diffuse arrangement and a subtle leftward bias in the directed syllables compared to the undirected syllables. Figure 4b displays the same syllables with respect to the first two principal components of the VAE’s latent features, showing a much more concentrated distribution of directed syllables relative to undirected syllables (see Figure S9 for all syllables). In fact, when we quantify this reduction of variability across all feature-space dimensions and song syllables (see Methods), learned latent features consistently report greater variability reductions than SAP-generated features (Figure 4c; SAP: 0-20%, VAE: 27-37%) indicating latent features are more sensitive to this effect. Additionally, we find that latent features outperform SAP features in the downstream task of predicting social context (Table S1).

**Figure 4:**
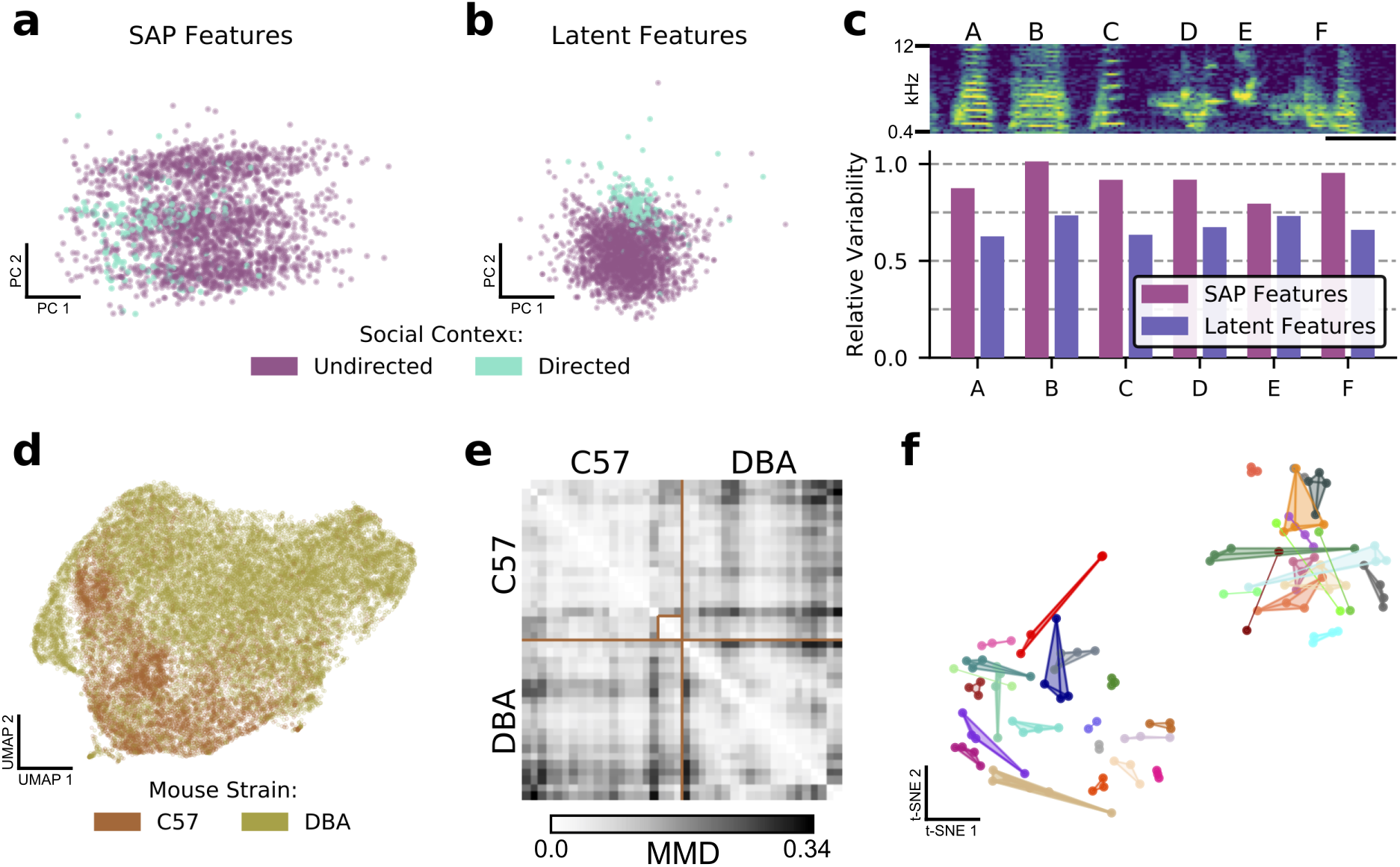
Latent features better capture differences in sets of vocalizations. **a)** The first two principal components in SAP feature space of a single zebra finch song syllable, showing differences in directed and undirected syllable distributions. **b)** The first two principal components of latent syllable features, showing the same comparison. Learned latent features more clearly indicate differences between the two conditions by clustering directed syllables together. **c)** Acoustic variability of each song syllable as measured by SAP features and latent features (see Methods). Latent features more clearly represent the constriction of variability in the directed context. Spectrogram scale bar denotes 100ms. **d)** A UMAP projection of the latent means of USV syllables from two strains of mice, showing clear differences in their vocal repertoires. **e)** Similarity matrix between syllable repertoires for each pair of the 40 recording sessions from **d**. Lighter values correspond to more similar syllable repertoires (lower Maximum Mean Discrepancy (MMD)). **f)** t-SNE representation of similarities between syllable repertoires, where distance metric is estimated MMD. The dataset, which is distinct from that represented in **d** and **e**, contains 36 individuals, 118 recording sessions, and 156,180 total syllables. Color indicates individual mice, and scatterpoints of the same color represent repertoires recorded on different days. Distances between points represent the similarity in vocal repertoires, with closer points more similar. We note that the major source of repertoire variability corresponds to genetic background, corresponding to the two distinct clusters (Figure S11). A smaller level of variability can be seen across individuals in the same clusters. Individual mice have repertoires with even less variability, indicated by the close proximity of most repertoires from each mouse.

Similarly, we can ask whether latent features are able to capture differences between groups of individuals. In [50], the authors compared USVs of 12 strains of mice using a clustering-based approach. Here, we perform an alternative version of this analysis using two mouse strains (C57/BL6 and DBA/2) from a publicly available dataset that were included in this earlier study. Figure 4d shows a UMAP projection of the 31,440 detected syllables, colored by mouse strain. Visualized with UMAP, clear differences between the USV distributions are apparent. While in contrast to traditional acoustic features such as ‘mean frequency,’ individual VAE latent features (vector components) are generally less interpretable, when taken together with an “atlas” of USV shapes derived from this visualization (Figure S10), we can develop an intuitive understanding of the differences between the USVs of the two strains: the C57 mice mostly produce noisy USVs, while the DBA mice produce a much greater variety, including many short low-frequency syllables that C57s rarely produce.

Given these results, we asked whether these strain differences are evident at the level of individual 6.5-minute recording sessions. To compare distributions of syllables without making restrictive parametric assumptions, we employed Maximum Mean Discrepancy (MMD), a difference measure between pairs of distributions [14]. We estimated MMD between the distributions of latent syllable encodings for each pair of recording sessions (see Methods) and visualized the result as a distance matrix (Figure 4e). Here, lighter values indicate more similar syllable repertoires. We note that, in general, values are brighter when comparing repertoires within strains than when comparing across strains, consistent with the hypothesis of inter-strain differences. We also note some substructure, including a well-defined cluster within the C57 block (annotated).

Finally, we used a much larger library of female-directed mouse USVs (36 individuals, 2-4 20-minute recording sessions each, 40 total hours of audio, 156,000 syllables) to investigate the diversity and stability of syllable repertoires. We repeated the above procedure, estimating MMD for each pair of recording sessions (Figure S11), and then computed a t-SNE layout of the recording sessions with estimated MMD as the distance metric to visualize the distribution of syllable repertoires (see Methods). In Figure 4f, each recording session is represented by a scatterpoint, and recordings of the same individual are connected and displayed in the same color. We note an overall organization of syllables into two clusters, corresponding to the genetic backgrounds of the mice (Figure S11). Furthermore, we note that almost all recordings of the same individuals are co-localized, indicating that within-subject differences in syllable repertoire are smaller than those between individuals. Although it has been previously shown that a deep convolutional neural network can be trained to classify USV syllables according to mouse identity with good accuracy ([19], Figure S1), here we find that repertoire features learned in a wholly unsupervised fashion achieve similar results, indicating mice produce individually-stereotyped, stable vocal repertoires.

### 2.4 Latent features fail to support cluster substructure in USVs

Above, we have shown that, by mapping complex sets of vocalizations to low-dimensional latent representations, VAEs allow us to visualize the relationships among elements in mouse vocal repertoires. The same is likewise true for songbirds such as the zebra finch, *T. guttata*. Figure 5 compares the geometry of learned latent spaces for an individual of each species as visualized via UMAP. As expected, the finch latent space exhibits well-delineated clusters corresponding to song syllables (Figure 5a). However, as seen above, mouse USVs clump together in a single quasi-continuous mass (Figure 5b). This raises a puzzle, since the clustering of mouse vocalizations is often considered well-established in the literature [18, 4, 53, 6, 16] and is assumed in most other analyses of these data [50, 7]. Clusters of mouse USVs are used to assess differences across strains [50], social contexts [6, 7, 15], and genotypes [13], and the study of transition models among clusters of syllables has given rise to models of syllable sequences that do not readily extend to the non-clustered case [18, 6, 16].

**Figure 5:**
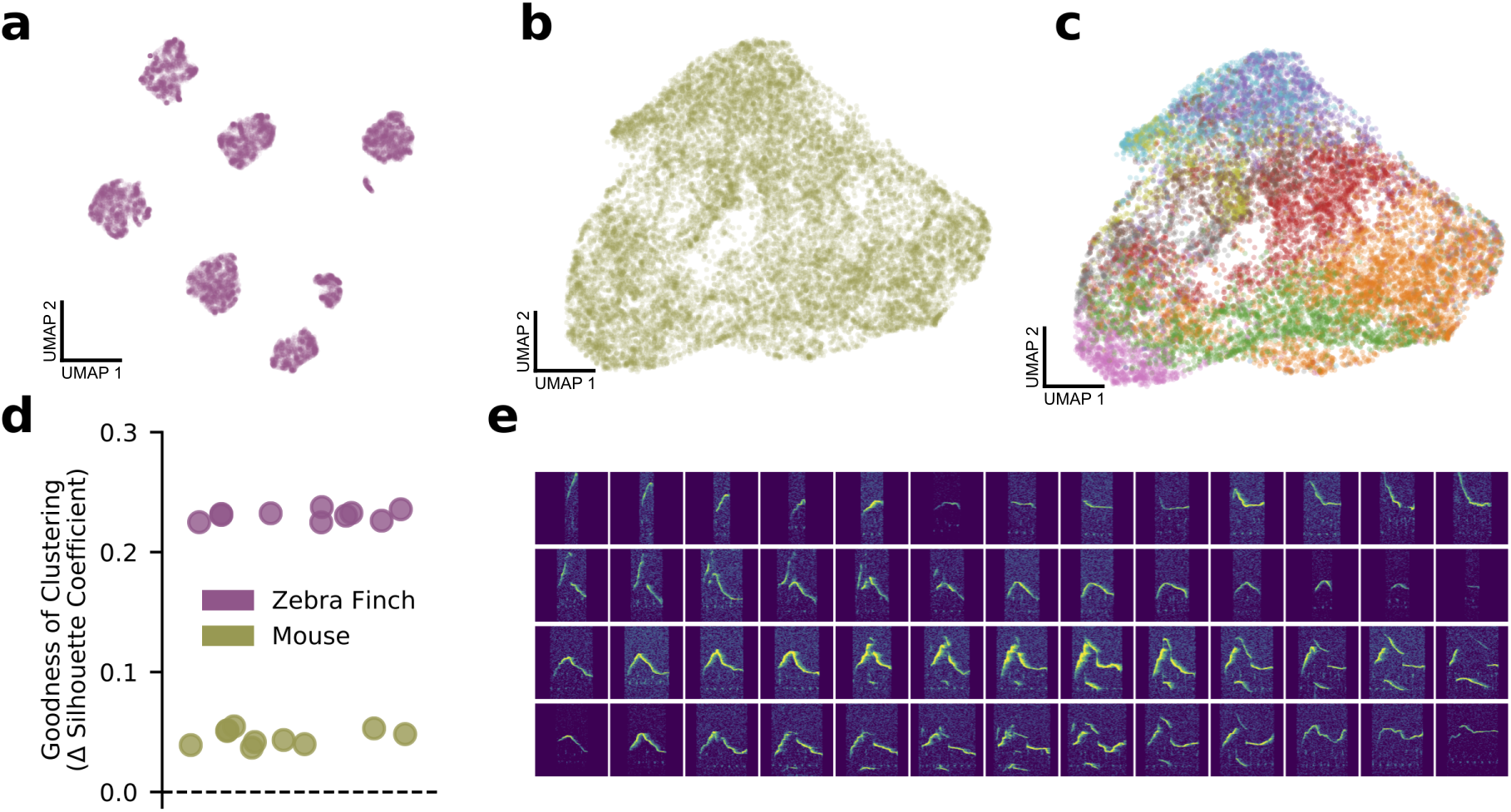
Bird syllables clearly cluster, but mouse USVs do not. **a)** UMAP projection of the song syllables of a single male zebra finch (14,270 syllables) **b)** UMAP projection of the USV syllables of a single male mouse (17,400 syllables) **c)** the same UMAP projection as in **b**, colored by MUPET-assigned labels **d)** Mean silhouette coefficient (an unsupervised clustering metric) for latent descriptions of zebra finch song syllables and mouse syllables. The dotted line indicates the null hypothesis of a single covariance-matched Gaussian noise cluster fit by the same algorithm. Each scatterpoint indicates a cross-validation fold, and scores are plotted as differences from the null model. Higher scores indicate more clustering. **e)** Interpolations (horizontal series) between distinct USV shapes (left and right edges) demonstrating the lack of data gaps between putative USV clusters.

We therefore asked whether mouse USVs do, in fact, cluster or whether, as the latent space projection suggests, they form a single continuum. In principle, this is impossible to answer definitively, because, without the benefit of ground truth labels, clustering is an unsupervised classification task. Moreover, there is little consensus among researchers as to the best method for assessing clustering and where the cutoff between clustered and non-clustered data lies [20]. In practice, new clustering algorithms are held to function well when they outperform previous approaches and produce sensible results on data widely agreed on to be clustered. Thus, while it is clear that zebra finch song syllables should be and are clustered by the VAE (Figure 5a), we can only ask whether clustering is a more or less satisfying account of the mouse data in Figure 5b.

To address this question, we performed a series of analyses to examine the clustering hypothesis from complementary angles. First, we asked how clusters detected by other analysis approaches correspond to regions in the latent space. As shown in Figure 5c, clusters detected by MUPET roughly correspond to regions of the UMAP projection, with some overlap between clusters (e.g., purple and blue clusters) and some non-contiguity of single clusters (red and orange clusters). That is, even though clusters do broadly label different subsets of syllables, they also appear to substantially bleed into one another, unlike the finch song syllables in Figure 5a. However, it might be objected that Figure 5b displays the UMAP projection, which only attempts to preserve local relationships between nearest neighbors and is not to be read as an accurate representation of the latent geometry. Might the lack of apparent clusters result from distortions produced by the projection to two dimensions? To test this, we calculated several unsupervised clustering metrics on full, unprojected latent descriptions of zebra finch and mouse syllables. By these measures, both bird syllables and mouse USVs were more clustered than moment-matched samples of Gaussian noise, a simple null hypothesis, but mouse USVs were closer to the null than to birdsong on multiple goodness-of-clustering metrics (Figure 5d, S12). Additionally, we find that MUPET acoustic features admit uniformly poorer clusters than latent features as quantified by the same metrics (Figure S13). On a more practical note, we compared the consistency of cluster labels assigned by Gaussian mixture models trained on disjoint subsets of zebra finch and mouse latent syllable descriptions to determine how well the structure of the data determines cluster membership across repeated fittings (Figure S15). We find near-perfectly consistent assignments of zebra finch syllables into six clusters and much less consistent clusters for mouse syllables for more than two clusters. Finally, we tested whether the data contained noticeable gaps between syllables in different clusters. If syllable clusters are well-defined, there should not exist smooth sequences of data points connecting distinct examples. However, we find that even the most acoustically disparate syllables can be connected with a sequence of syllables exhibiting more-or-less smooth acoustic variation (5e), in contrast to zebra finch syllables (Figure S16). Thus, even though clustering may not constitute the best account of mouse USV syllable structure, learned latent features provide useful tools to both explore and quantify the acoustic variation within and across species.

### 2.5 Measuring acoustic variability over tens of milliseconds

The results above have shown that data-derived latent features represent more information about syllables than traditional metrics and can successfully capture differences within and between individuals and groups. Here, we consider how a related approach can also shed light on the short-time substructure of vocal behavior.

The analysis of syllables and other discrete segments of time is limited in at least two ways. First, timing information, such as the lengths of gaps between syllables, is ignored. Second, experimenters must choose the unit of analysis (syllable, song motif, bout), which has a significant impact on the sorts of structure that can be identified [22]. In an attempt to avoid these limitations, we pursued a complementary approach, using the VAE to infer latent descriptions of fixed duration audio segments, irrespective of syllable boundaries. Similar to the shotgun approach to gene sequencing [51] and a related method of neural connectivity inference [47], we trained the VAE on randomly sampled segments of audio, requiring that it learn latent descriptions sufficient to characterize any given time window during the recording. That is, this “shotgun-VAE” approach encouraged the autoencoder to find latent features sufficient to “glue” continuous sequences back together from randomly sampled audio snippets.

Figure 6a shows a UMAP projection of latent features inferred from fixed-duration segments from a subset of the mouse USVs shown in Figure 5b. While this projection does display some structure (silence on the right, shorter to longer syllables arranged from right to left), there is no evidence of stereotyped sequential structure. In contrast, Figure 6b shows the same technique applied to bouts of zebra finch song, with the song represented as a single well-defined strand coursing clockwise from the bottom to the top left of the projection. Other notable features are the loop on the left containing repeated, highly variable introductory notes that precede and often join song renditions and a “linking note” that sometimes joins song motifs. Most importantly, such a view of the data clearly illustrates not only stereotypy but variability: introductory notes are highly variable, but so are particular syllables (B, E) in contrast to others (C, F).

**Figure 6:**
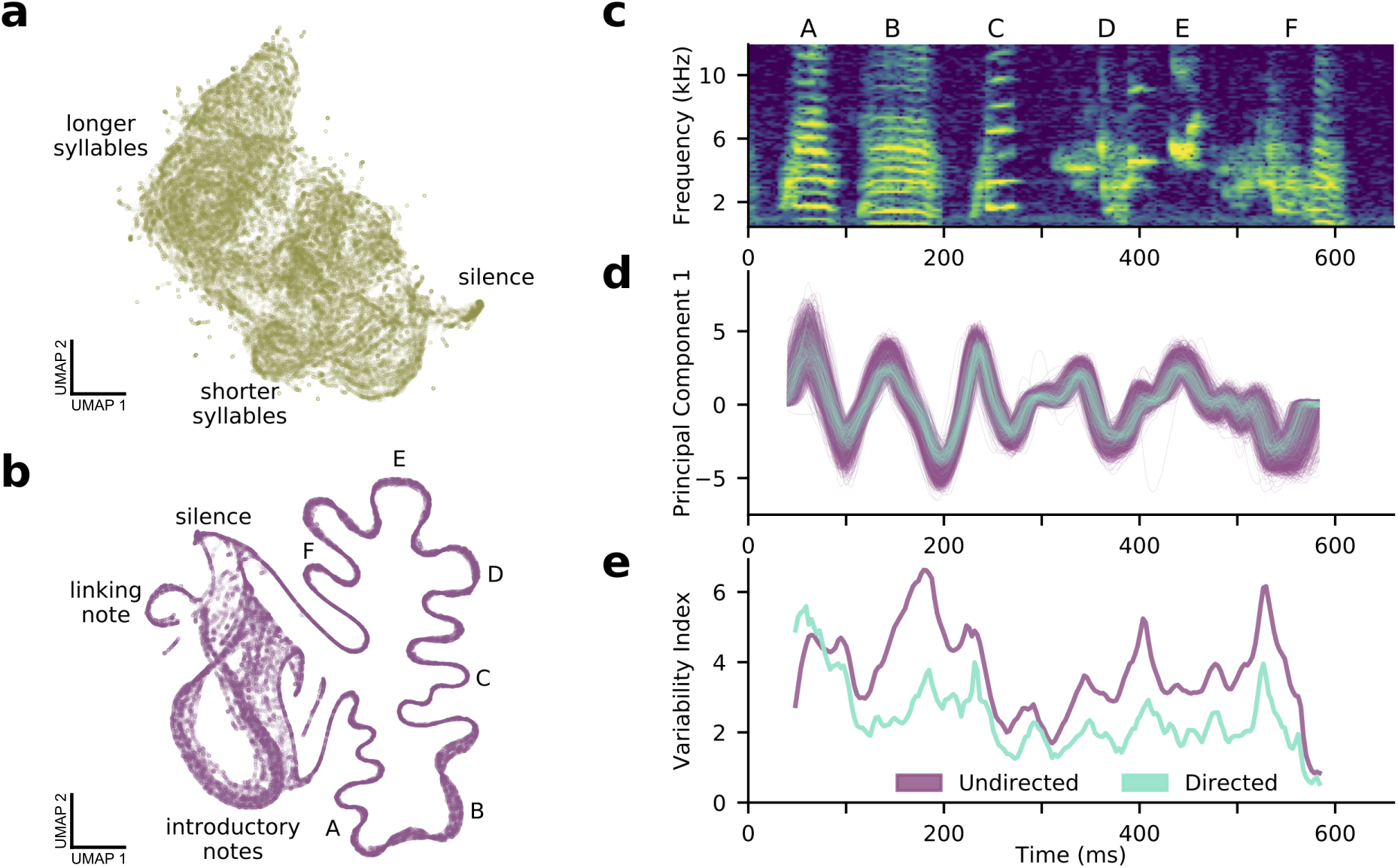
A shotgun VAE approach learns low dimensional latent representations of subsampled, fixed-duration spectrograms and captures short-timescale variability in behavior. **a)** A UMAP projection of 100,000 200ms windows of mouse USVs (cp. Figure 4a). **b)** A UMAP projection of 100,000 120ms windows of zebra finch song (cp. Figure 4b). Song progresses counterclockwise on the right side, while more variable, repeated introductory notes form a loop on the left side. **c)** A single rendition of the song in **b**. **d)** The song’s first principal component in latent space, showing both directed (cyan) and undirected (purple) renditions. **e)** In contrast to a syllable-level analysis, the shotgun approach can measure zebra finch song variability in continuous time. Song variability in both directed (cyan) and undirected (purple) contexts are plotted (see Methods).

Following this, we asked whether the shotgun VAE method could be used to assess the phenomenon of reduced variability in directed birdsong [46]. We examined the song portion of Figure 6b in both directed and undirected conditions, warping each in time to account for well-documented differences in rhythm and tempo. We then trained a VAE on randomly sampled 80ms portions of the warped spectrograms. As a plot of the first principle component of the latent space shows (Fig. 4d), the VAE is able to recover the expected reduction in directed song variability on a tens-of-milliseconds timescale relevant to the hypothesized neural underpinnings of the effect [11]. This result recapitulates similar analyses that have focused on harmonic and tonal syllables like A and B in Figure 4c [21], but the shotgun VAE method is applicable to all syllables, yielding a continuous estimate of song variability (Fig. 4e). Thus, not only do VAE-derived latent features capture structural properties of syllable repertoires, the shotgun VAE approach serves to characterize continuous vocal dynamics as well.

Video 1: An animated version of Figure 6a. Recorded USVs from a single male mouse are played while the corresponding latent features are visualized by a moving star in a UMAP projection of latent space. The recording is slowed by a factor of four and additionally pitch shifted downward by a factor of two so that the vocalizations are audible.

Video 2: An animated version of Figure 6b. A recorded song bout from a single male zebra finch is played while the corresponding latent features are visualized by a moving star in a UMAP projection of latent space.

### 2.6 Latent features capture song similarity

Above, we saw how a subsampling-based “shotgun VAE” approach can capture fine details of zebra finch vocal behavior modulated by social context. However, a principal reason songbirds are studied is their astonishing ability to copy song. A young male zebra finch can successfully learn to sing the song of an older male over the course of multiple months after hearing only a handful of song renditions. At least three methods exist for quantifying the quality of song learning outcomes, with two using hand-picked acoustic features [49, 48, 30] and another using Gaussian distributions in a similarity space based on power spectral densities to represent syllable categories [32]. We reiterate that hand-picked acoustic features are sensitive to experimenter choices, with some acoustic features like pitch only defined for certain kinds of sounds. Additionally, restrictive parametric assumptions limit the potential sensitivity of a method. By contrast, the VAE learns a compact feature representation of complete spectrograms and MMD provides a convenient nonparametric difference measure between distributions. Therefore, as a final assessment of the VAE’s learned features, we asked whether latent features reflect the similarity of tutor and pupil songs.

To assess the VAE’s ability to capture effects of song learning, we trained a syllable VAE and a shotgun VAE on song motifs of ten paired adult zebra finch tutors and adult zebra finch pupils. As Figure 7a shows for the syllable-level analysis, most syllables form distinct clusters in a latent UMAP embedding, with many tutor/pupil syllable pairs sharing a cluster. For a specific tutor/pupil pair shown in Figure 7b, we highlight three corresponding syllables from the two motifs. The first and third syllables (C and E) are well-copied from tutor to pupil, but the pupil’s copy of the second syllable (syllable D) does not contain the high frequency power present in the first half of the tutor’s syllable. This discrepancy is reflected in the latent embedding, with the two versions of syllable D corresponding to distinct clusters. We quantified the quality of copying by calculating maximum mean discrepancy (MMD) between each pair of tutor and pupil syllables, shown in 7c. A dark band of low MMD values along the diagonal indicates well-copied syllables and syllable orderings for most birds.

**Figure 7:**
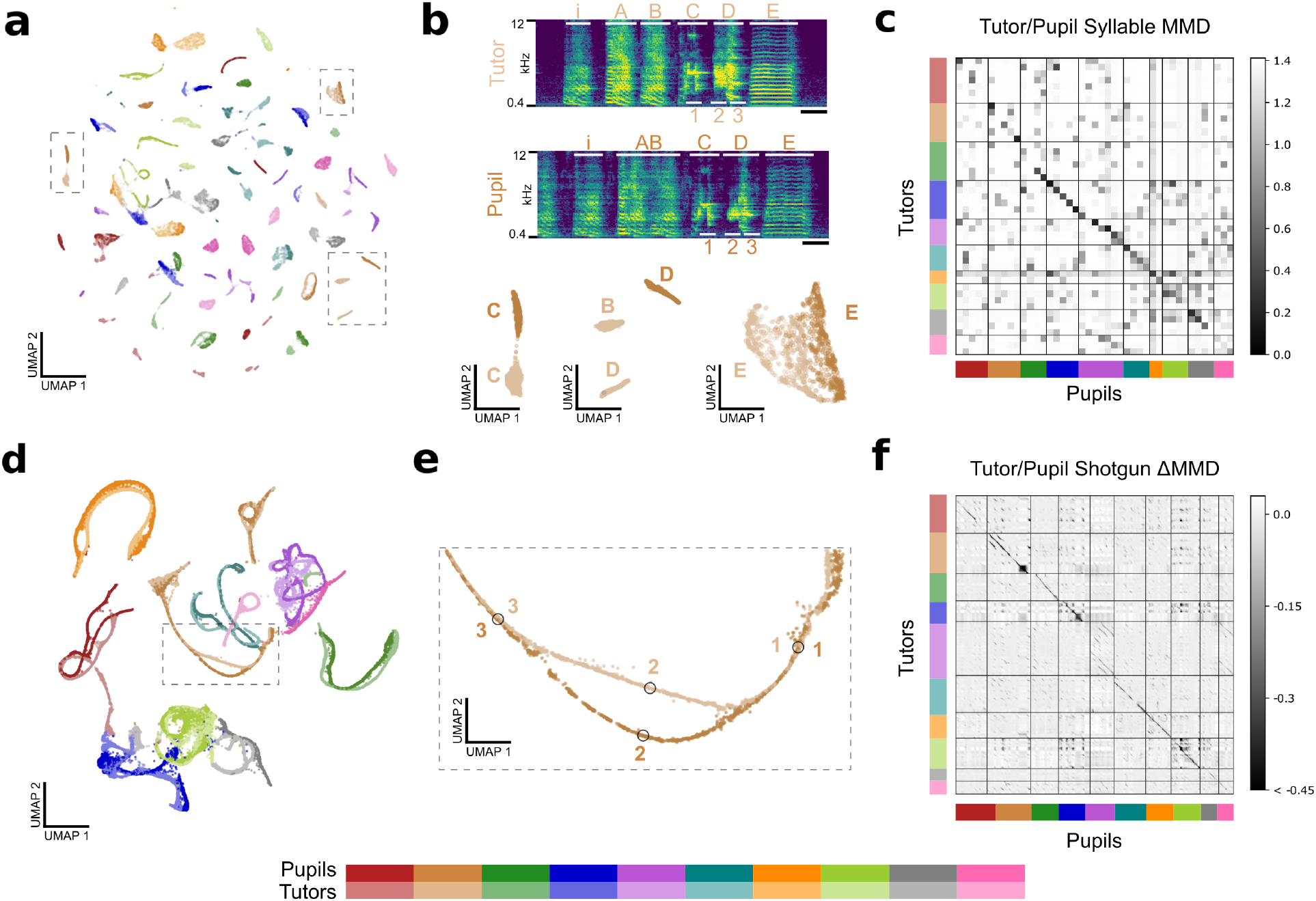
Latent features capture similarities between tutor and pupil song. **a)** Latent UMAP projection of the song syllables of 10 zebra finch pupil/tutor pairs. Note that many tutor and pupil syllables cluster together, indicating song copying. **b)** Example song motifs from one tutor/pupil pair. Syllables “C” and “E” are well-copied, but the pupil’s rendition of syllable “D” does not have as much high-frequency power as the tutor’s rendition. The difference between these two renditions is captured in the latent UMAP projection below. Additionally, the pupil sings a concatenated version of the tutor’s syllables “A” and “B,” a regularity that a syllable-level analysis cannot capture. Thus, the pupil’s syllable “AB” does not appear near tutor syllable “B” in the UMAP projection. Scale bar denotes 100ms. **c)** Quality of song copying, estimated by maximum mean discrepancy (MMD) between every pair of tutor and pupil syllables. The dark band of low MMD values near the diagonal indicates good song copying. Syllables are shown in motif order. **d)** Latent UMAP projection of shotgun VAE latents (60ms windows) for the song motifs of 10 zebra finch pupil/tutor pairs. Song copying is captured by the extent to which pupil and tutor strands co-localize. **e)** A detail of the UMAP projection in panel **d** shows a temporary split in pupil and tutor song strands spanning the beginning of syllable “D” with poorly copied high-frequency power. Labeled points correspond to the motif fragments marked in panel **b**. **f)** MMD between pupil and tutor shotgun VAE latents indexed by continuous-valued time-in-motif quantifies song copying on fine timescales. The dark bands near the diagonal indicate well-copied stretches of song. The deviation of MMD values from a rank-one matrix is displayed for visual clarity (see Figure S21 for details). Best viewed zoomed in.

To complement the previous analysis, and to sidestep difficulties related to syllable segmenting such as the fused “AB” syllable learned by the pupil in Figure 7b, we performed an analogous analysis using the shotgun VAE approach. After training the VAE on 60ms chunks of audio drawn randomly from each song motif, we projected the learned latent features into two dimensions using UMAP (Figure 7d) with a modified distance to prevent motif “strands” from breaking (see Methods, Figure S19). As expected, we see a close correspondence between pupil and tutor motif strands, with only a single pupil/tutor pair (pink) not overlapping. In fact, out of the roughly 3.6 million possible pairings of pupils and tutors, it is easily apparent which is the true pairing. Additionally, we find that the finer details are informative. For instance, the poorly copied syllable “D” from Figure 7b corresponds to a temporary divergence of the pupil strand from the tutor strand (Figure 7e). We quantified the quality of copying in continuous time by calculating MMD between the distributions of song latents corresponding to distinct times within motifs. Deviations from a simple rank-one structure are shown in Figure 7f (see Figure S21 for the original MMD matrix). Consistent with Figure 7c, a dark band near the diagonal indicates good copying, quantifying the quality of copying in much more detail than a syllable-level analysis could provide.

Video 3: An animated version of Figure 7d. Recorded songs motifs from each male zebra finch is played while the corresponding latent features are visualized by a moving star in a UMAP projection of latent space.

## Discussion

The complexity and high dimensionality of vocal behavior have posed a persistent challenge to the scientific study of animal vocalization. In particular, comparisons of vocalizations across time, individuals, groups, and experimental conditions require some means of characterizing the similarity of selected groups of vocal behaviors. Feature vector-based approaches and widespread software tools have gone a long way toward addressing this challenge and providing meaningful scientific insights, but the reliance of these methods on handpicked features leaves open the question of whether other feature sets might better characterize vocal behavior.

Here, we adopt a data-driven approach, demonstrating that features learned by the variational autoencoder (VAE), an unsupervised learning method, outperform frequently used acoustic features across a variety of common analysis tasks. As we have shown, these learned features are both more parsimonious (Figure S2), capture more variability in the data (Figure 2e,f), and better characterize vocalizations as judged by nearest neighbor similarity (Figure 3, S6, S7). Moreover, these features easily facilitate comparisons across sessions (Figure 4f), social contexts (Figure 4a-c), and individuals (Figure 4d-f, 7), quantifying not only differences in mean vocal behavior (Figure 4d), but also in vocal variability (Figure 4c).

This data-driven approach is closely related to previous studies that have applied dimensionality reduction algorithms (UMAP [31] and t-SNE [29]) to spectrograms to aid in syllable clustering of birdsong [42] and to visualize juvenile song learning in the zebra finch [26]. Additionally, a related recent publication [43] similarly described the application of UMAP to vocalizations of several more species and the application of the VAE to generate interpolations between birdsong syllables for use in playback experiments. Here, by contrast, we restrict use of the UMAP and t-SNE dimensionality reduction algorithms to visualizing latent spaces inferred by the VAE and use the VAE as a general-purpose tool for quantifying vocal behavior, with a focus on cross-species comparisons and assessing variability across groups, individuals, and experimental conditions.

Moreover, we have argued above that, despite conventional wisdom, clustering is not the best account of the diversity of mouse vocal behavior. We argued this on the basis of multiple converging lines of evidence, but note three important qualifications: First, the huge variety of vocal behavior among rodents [2, 18, 35, 41, 44, 33] suggests the possibility of clustered vocal behavior in some mouse strains not included in our data. Second, it is possible that the difference in clustered and non-clustered data depends crucially on data set size. If real syllables even occasionally fall between well-defined clusters, a large enough data set might lightly “fill in” true gaps. Conversely, even highly clustered data may look more or less continuous given an insufficient number of samples per cluster. While this is not likely given the more than 15,000 syllables in Figure 5, it is difficult to rule out in general. Finally, our purely signal-level analysis of vocal behavior cannot address the possibility that a continuous distribution of syllables could nevertheless be perceived categorically. For example, swamp sparrows exhibit categorical neural and behavioral responses to changes in syllable duration [38]. Nonetheless, we argue that, without empirical evidence to this effect in rodents, caution is in order when interpreting the apparent continuum of USV syllables in categorical terms.

Lastly, we showed how a “shotgun VAE” approach can be used to extend our approach to the quantification of moment-by-moment vocal variability. In previous studies, syllable variability has only been quantified for certain well-characterized syllables like harmonic stacks in zebra finch song [21]. Our method, by contrast, provides a continuous variability measure for all syllables (Figure 6c). This is particularly useful for studies of the neural basis of this vocal variability, which is hypothesized to operate on millisecond to tens-of-milliseconds timescales [11].

Nonetheless, as a data-driven method, our approach carries some drawbacks. Most notably, the VAE must be trained on a per-dataset basis. This is more computationally intensive than calculating typical acoustic features (≈1 hour training times on a GPU) and also prevents direct comparisons across datasets unless they are trained together in a single model. Additionally, the resulting learned features, representing nonlinear, non-separable acoustic effects, are somewhat less interpretable than named acoustic features like duration and spectral entropy. However, several recent studies in the VAE literature have attempted to address this issue by focusing on the introduction of covariates [45, 28, 23] and “disentangling” approaches that attempt to learn independent sources of variation in the data [17, 3], which we consider to be promising future directions. We also note that great progress in generating raw audio has been made in the past few years, potentially enabling similar approaches that bypass an intermediate spectrogram representation [36, 27].

Finally, we note that while our focus in this work is vocal behavior, our training data are simply syllable spectrogram images. Similar VAE approaches could also be applied to other kinds of data summarizable as images or vectors. The shotgun VAE approach could likewise be applied to sequences of such vectors, potentially revealing structures like those in Figure 6b. More broadly, our results suggest that data-driven dimensionality reduction methods, particularly modern nonlinear, overparameterized methods, and the latent spaces that come with them, offer a promising avenue for the study of many types of complex behavior.

## 3 Methods

### Animal Statement

All experiments were conducted according to protocols approved by the Duke University Institutional Animal Care and Use Committee.

### Recordings

Recordings of C57BL/6 and DBA/2 mice were obtained from the MUPET Wiki [1]. These recordings are used in Figures 2, 3, 4d-e, and 5e, S1, S2, S3, S4, S5, S6, S7, S8, and S10 and Table S2.

Additional recordings of female-directed mouse USVs are used in Figures 4f and S11. These recordings comprise 36 male mice from various genetic backgrounds over 118 recording sessions of roughly 20 minutes each (≈40 total hours, 156,180 total syllables). USVs were recorded with an ultrasonic microphone (Avisoft, CMPA/CM16), amplified (Presonus TubePreV2), and digitized at 300 kHz (Spike 7, CED). A subset of these recordings corresponding to a single individual (17,400 syllables) are further used in Figure 5b-d, 6a, S12, S13, S15, S17, and S20. Because these recordings contained more noise than the first set of C57/DBA recordings, we removed false positive syllables by training the VAE on all detected syllables, projecting latent syllables to two dimensions, and then removing syllables contained within the resulting cluster of noise syllables, with 15,712 syllables remaining (see Figure S20).

A single male zebra finch was recorded over two-day period (153-154 days post-hatch) in both female-directed and undirected contexts (14,270 total syllables, 1100 directed). Song was recorded with Sound Analysis Pro 2011.104 [48] in a soundproof box. These recordings are used in Figures 2, 4, 5, 6, S1, S2, S3, S4, S5, S7, S9, S12, S14, S15, S16, S17, and S18 and Table S1.

For Figure 7, we selected 10 adult, normally reared birds from different breeding cages in our colony. Until at least 60 days-post-hatch (dph), each of these pupil birds had interacted with only one adult male, the tutor from his respective breeding cage. We recorded the adult (>90 dph) vocalizations of these pupil birds for 5 to 12 days each with Sound Analysis Pro 2011.104 [48], then recorded their respective tutors under the same conditions for 5 to 12 days each. These recordings are used in Figures 7 and S19.

### Software Comparisons

We compared our VAE method to three widely used vocal analysis packages: MUPET 2.0 [50], DeepSqueak 2.0 [7] (for mouse USVs) and SAP 2011.104 [48] (for birdsong), each with default parameter settings. MUPET clusters were found using the minimum number of clusters (10). DeepSqueak features were generated using the DeepSqueak “import from MUPET” feature.

### Audio Segmenting

For all mouse USV datasets, individual syllable onsets and offsets were detected using MUPET with default parameter settings. The shotgun VAE analysis in Figure 6 was restricted to manually-defined regions (bouts) of vocalization. In this figure, zebra finch songs were segmented semi-automatically: first we selected four representative song motifs from each individual. Then we converted these to spectrograms using a Short Time Fourier Transform with Hann windows of length 512 and overlap of 256, averaged these spectrograms, and blurred the result using a gaussian filter with 0.5 pixel standard deviation. The result was a song template used to match against the remaining data. Specifically, we looked for local maxima in the normalized cross-correlation between the template and each audio file. Matches corresponded to local maxima with cross-correlations above 1.8 median absolute deviations from the median, calculated on a per-audio-file basis. A spectrogram was then computed for each match. All match spectrograms were then projected to two dimensions using UMAP [31], from which a single well-defined cluster, containing stereotyped song, was retained. Zebra finch syllable onsets and offsets were then detected using SAP [48] on this collection of song renditions using default parameter settings. After segmentation, syllable spectrograms were projected to two dimensions using UMAP, and eight well-defined clusters of incorrectly segmented syllables were removed, leaving six well-defined clusters of song syllables. For Figure 7, song motifs were hand-labeled for approximately 10 minutes of song-rich audio per animal. These labels were used to train an automated segmentation tool, *vak* 0.3.1 [34], for each animal. Trained *vak* models were used to automatically label motifs in the remaining audio data for each animal. Automatic segmentation sometimes divided single motifs or joined multiple motifs. To correct for these errors, short (<50 ms) gaps inside motifs were eliminated. After this correction, putative motif segments with durations outside 0.4s to 1.5s were discarded. Syllables segments were derived from a subset of the *vak* motif segments by aligning the motif amplitude traces and manually determining syllable boundaries, resulting in 75,430 total syllable segments.

### Spectrograms

Spectrograms were computed using the log modulus of a signal’s Short Time Fourier Transform, computed using Hann windows of length 512 and overlap of 256 for bird vocalization, and length 1024 and overlap 512 for mouse vocalization. Sample rates were 32kHz for bird vocalization and 250kHz for mouse vocalization, except for the recordings in Figure 3f, which were sampled at a rate of 300kHz. The resulting time/frequency representation was then interpolated at desired frequencies and times. Frequencies were mel-spaced from 0.4 to 8kHz for bird recordings in Figure 7, mel-spaced from 0.4 to 10kHz for all other bird recordings, and linearly spaced from 30 to 110kHz for mouse recordings. For both species, syllables longer than *t*_max_ = 200ms were discarded. Additionally, short syllables were stretched in time in a way that preserved relative duration, but encouraged the VAE to represent fine temporal details. Specifically, a syllable of duration *t* was stretched by a factor of 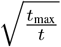. See Figure S4 for a comparison to linear frequency spacing for zebra finches and no time stretching for mice and zebra finches. The resulting spectrograms were then clipped to manually tuned minimum and maximum values. The values were then linearly stretched to lie in the interval [0, 1]. The resulting spectrograms were 128 *×* 128 = 16384 pixels, with syllables shorter than *t*_max_ zero-padded symmetrically.

### Model Training

Our variational autoencoder is implemented in PyTorch (v1.1.0) and trained to maximize the standard evidence lower bound (ELBO) objective using the reparameterization trick and ADAM optimization [25, 39, 37, 24]. The encoder and decoder are deep convolutional neural networks with fixed architecture diagrammed in Figure S22. The latent dimension was fixed to 32, which was found to be sufficient for all training runs. The approximate posterior was parameterized as a normal distribution with low rank plus diagonal covariance: 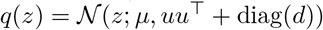 where *μ* is the latent mean, *u* is a 32×1 covariance factor, and *d* was the latent diagonal, a vector of length 32. The observation distribution was parameterized as 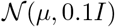 where *μ* was the output of the decoder. All activation functions were Rectified Linear Units. Learning rate was set to 10^−3^ and batch size was set to 64.

### Comparison of VAE and handpicked features

For each analysis tool (MUPET, DeepSqueak, SAP), we assembled two feature sets: one calculated by the comparison tool (e.g., MUPET featurs) and one a matched VAE set. For the first set, each feature calculated by the program was z-scored and all components with non-zero variance were retained (9/9, 10/10, and 13/14 components for MUPET, DeepSqueak, and SAP, respectively). For the second set, we trained a VAE on all syllables, computed latent means of these via the VAE encoder, and removed principal components containing less than 1% of the total feature variance (7, 5, and 5 out of 32 components retained for MUPET, DeepSqueak, and SAP syllables, respectively). Each feature set was used as a basis for predicting the features in the other set using *k*-nearest neighbors regression with *k* set to 10 and nearest neighbors determined using Euclidean distance in the assembled feature spaces. The variance-explained value reported is the average over 5 shuffled train/test folds (Figure 2e).

Unlike latent features, traditional features do not come equipped with a natural scaling. For this reason, we z-scored traditional features to avoid tethering our analyses to the identities of particular acoustic features involved. Then, to fairly compare the effective dimensionalities of traditional and acoustic features in Figure 2d, we thus also z-scored the latent features as well, thereby disregarding the natural scaling of the latent features. PCA was then performed on the resulting scaled feature set.

### Birdsong Variability Index

For Figure 4c, given a set {*z*_*i*_|*i* = 1 … *n*} of feature vectors of *n* syllables, we defined a variability index for the data as follows:

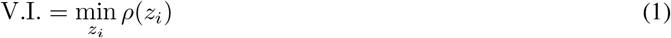

where *ρ*(*z*) is proportional to a robust estimator of the variance of the data around *z*:

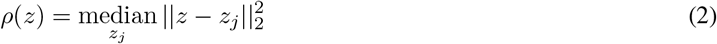

We calculate above metric for every combination of syllable (A-F), feature set (SAP-generated vs. VAE-generated), and social context (directed vs. undirected) and report the variability index of the directed condition relative to the variability index of the undirected condition (Figure 4c).

For Figure 6e, we would ideally use the variability index defined above, but *ρ*(*z*) is expensive to compute for each data point, as required in (1). Thus, we use an approximate center point defined by the median along each *coordinate*: 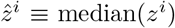, where the superscript here represents the *i*th coordinate of *z*. That is, 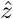 contains the medians of the marginal distributions. This value is calculated for each combination of timepoint and social context (directed vs. undirected) and plotted in Figure 6e.

### Maximum Mean Discrepancy

We used the Maximum Mean Discrepancy (MMD) integral probability metric to quantify differences in sets of syllables [14]. Given random variables *x* and *y*, MMD is defined over a function class 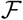 as 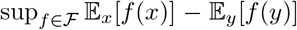. Here, 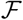 was taken to the set of functions on the unit ball in a reproducing kernel Hilbert space with fixed spherical Gaussian kernel. For Figures 4e-f, the kernel width *σ* was chosen to be the median distance between points in the aggregate sample, a common heuristic [14]. In Figure 7, the kernel bandwidth was chosen to be 25% of the median distance between points in the aggregate sample in order to focus on finer differences in distributions. For Figure 4e, we obtained 20 approximately 6.5 minute recordings of male C57BL/6 mice and 20 approximately 6.5 minute recordings of male DBA/2 mice (see Recordings). Latent means of USVs from a single recording were treated as independent and identically distributed draws from a recording-specific USV distribution, and MMD was estimated using these latent means. In Figure 4e, the MMD values are plotted as a matrix and the order of rows was obtained by agglomerative clustering. In Figures 4f and 7, the same procedure was followed. For Figure 4f, a t-distributed Stochastic Neighbor Embedding (t-SNE) was computed for each recording session, with the distance between recording sessions taken to be the estimated MMD between them (see Figure S11 for the MMD matrix).

### Unsupervised Clustering Metrics

We used three unsupervised clustering metrics to assess the quality of clustering for both zebra finch and mouse syllables: the mean silhouette coefficient [40], the Calinski-Harabasz Index [5], and the Davies-Bouldin Index [9]. For each species (zebra finch and mouse) we partitioned the data for tenfold cross-validation (train on 9/10, test on 1/10 held out). For a null comparison, for each of 10% disjoint subsets of the data, we created a synthetic Gaussian noise dataset matched for covariance and number of samples. These synthetic noise data sets were then used to produce the dotted line in Figure 5d.

For each data split, we clustered using a Gaussian Mixture Model (GMM) with full covariance using Expectation Maximization on the training set. We then evaluated each clustering metric on the test set. The number of clusters, *k*, was set to 6 in Figure 5d, but qualitatively similar results were obtained when *k* was allowed to vary between 2 and 12 (Figure S12). Reported values in Figure 5d and Figure S12 are the differences in unsupervised metrics on real data and Gaussian noise for each cross-validation fold, with a possible sign change to indicate higher values as more clustered.

### Shotgun VAE

To perform the analysis in Figure 6a-b, regions of active vocalization were defined manually for both species (22 minutes of mouse recordings, 2 minutes of zebra finch recordings). Zebra finch bouts containing only calls and no song motifs were excluded. For both species, the duration of audio chunks was chosen to be roughly as long as the longest syllables (zebra finch: 120ms; mouse: 200ms). No explicit training set was made. Rather, onsets and offsets were drawn uniformly at random from the set of fixed-duration segments and the corresponding spectrograms were computed on a per-datapoint basis. Thus, the VAE likely never encountered the same spectrogram twice, encouraging it to learn the underlying time series.

To perform the variability reduction analysis in Figure 6d-e, song renditions were collected (see Audio Segmenting) and a spectrogram was computed for each. The whole collection of spectrograms was then jointly warped using piecewise-linear time warping [52]. Fixed-duration training spectrograms were made by interpolating normal spectrograms (as described in Spectrograms) at linearly spaced time points in warped time, generally corresponding to non-linearly spaced time points in real time. As above, spectrograms were made during training on a per-datapoint basis. After training the VAE on these spectrograms, latent means were collected for 200 spectrograms for each song rendition, linearly spaced in warped time from the the beginning to the end of the song bout. Lastly, for each combination of condition (directed vs. undirected song) and timepoint, the variability index described above was calculated. A total of 186 directed and 2227 undirected song renditions were collected and analyzed.

To generate the shotgun VAE training set for Figure 7, 2000 *vak*-labeled motifs were selected from each animal (see Audio Segmenting). A single 60ms window was drawn from each motif to create a training set of 40,000 total spectrogram windows drawn from the 20-animal cohort. After training a VAE on this dataset, the hand-labeled motif segments used to train *vak* models (see Audio Segmenting) were segmented into overlapping 60ms windows that spanned each motif with an 8ms step size between successive windows (52,826 total windows). These windows were reduced with the trained VAE and their latent means subsequently analyzed.

### Modified UMAP

Although the standard UMAP embedding of shotgun VAE latents from single-finch datasets generates points along smoothly varying strands (see Figure 6b), UMAP typically broke motifs into multiple strand-like pieces in the 20-animal dataset from Figure 7. To encourage embeddings that preserve the neighbor relationship of successive windows, we modified the distance measure underlying the UMAP. First, we computed the complete pairwise Euclidean distance matrix between all windows in latent space. Then, we artificially decreased the distance between successive windows from the same motif by multiplying corresponding distance matrix entries by 10^−3^. This precomputed distance matrix was then passed to UMAP as a parameter. See Figure S19 for a comparison of the two UMAP projections.

## Data and Code Availability Statement

The latest version of Autoencoded Vocal Analysis, the Python package used to generate, plot, and analyze latent features, is freely available online: https://github.com/pearsonlab/autoencoded-vocal-analysis. Mouse and zebra finch recordings will be made available upon publication.

## 4 Supplementary Figures

**Figure S1:**
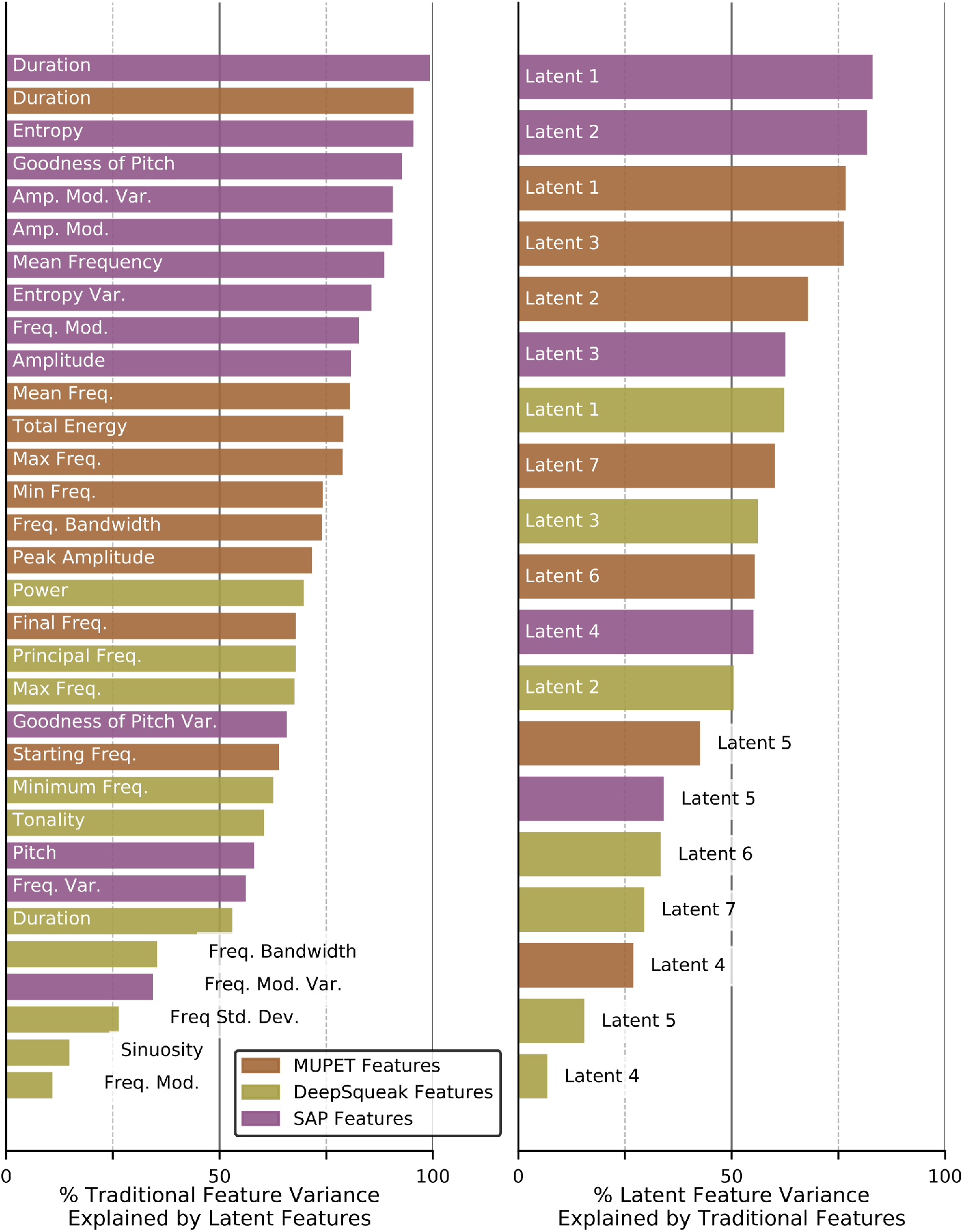
Left column: Named acoustic feature variance explained by latent features. Right column: Latent acoustic feature variance explained by named acoustic features. Values reported are the results of *k*-nearest neighbor classification averaged over five shuffled test/train folds (see Methods).

**Figure S2:**
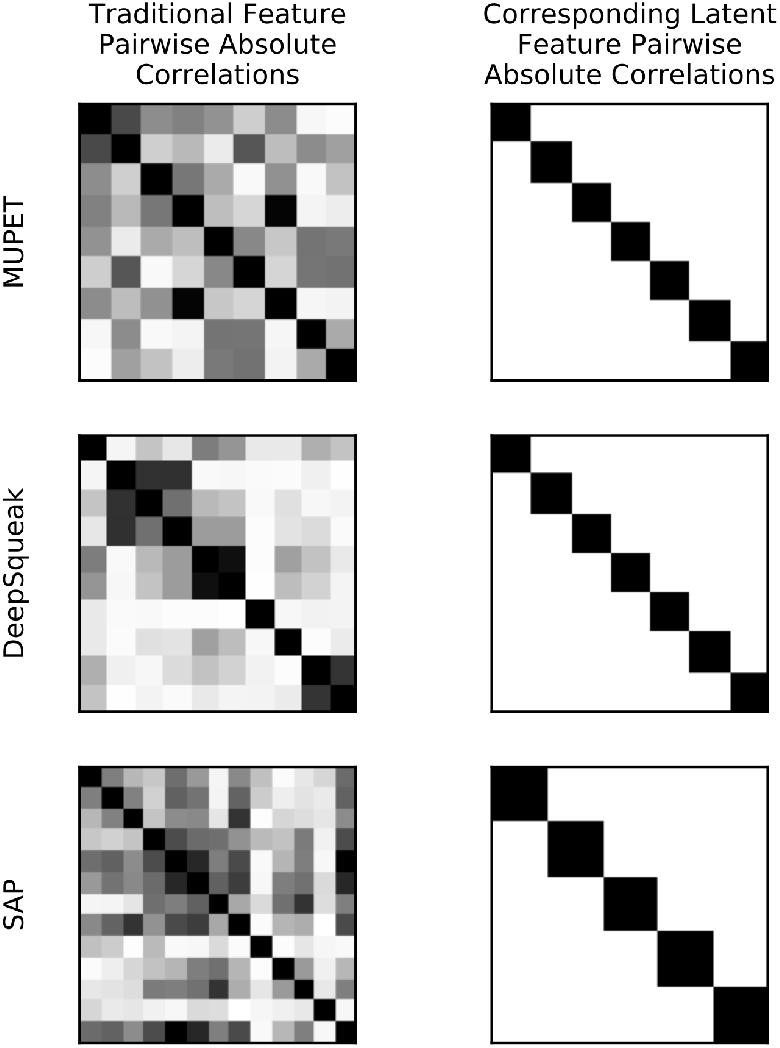
Traditional acoustic features are highly correlated. Left column: pairwise absolute correlations between named acoustic features when applied to the datasets in Figure 2. Right column: pairwise absolute correlations of latent features for the same datasets.

**Figure S3:**
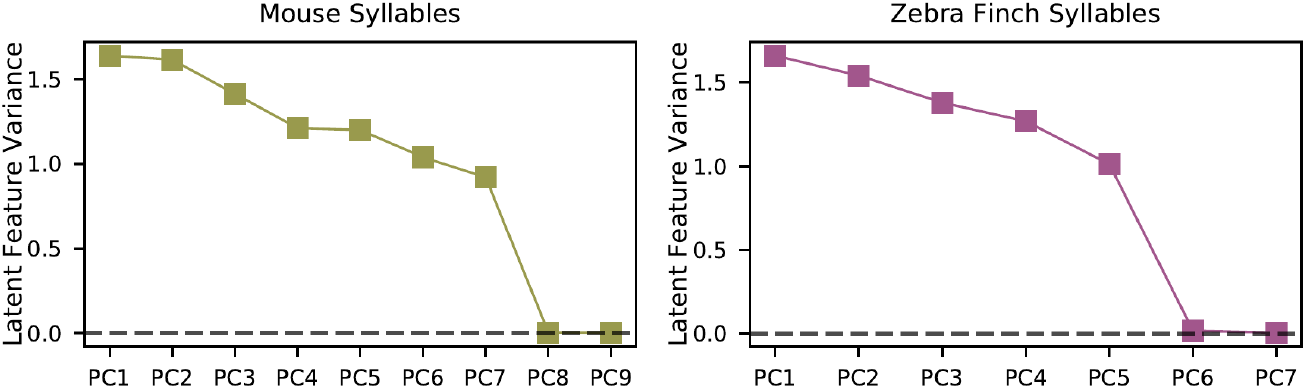
The VAE learns a parsimonious set of acoustic features. When trained on mouse syllables (from Figure 2a), the VAE makes use of only 7 of 32 latent dimensions. When trained on zebra finch syllables (from Figure 5a), the VAE makes use of only 5 of 32 latent dimensions.

**Figure S4:**
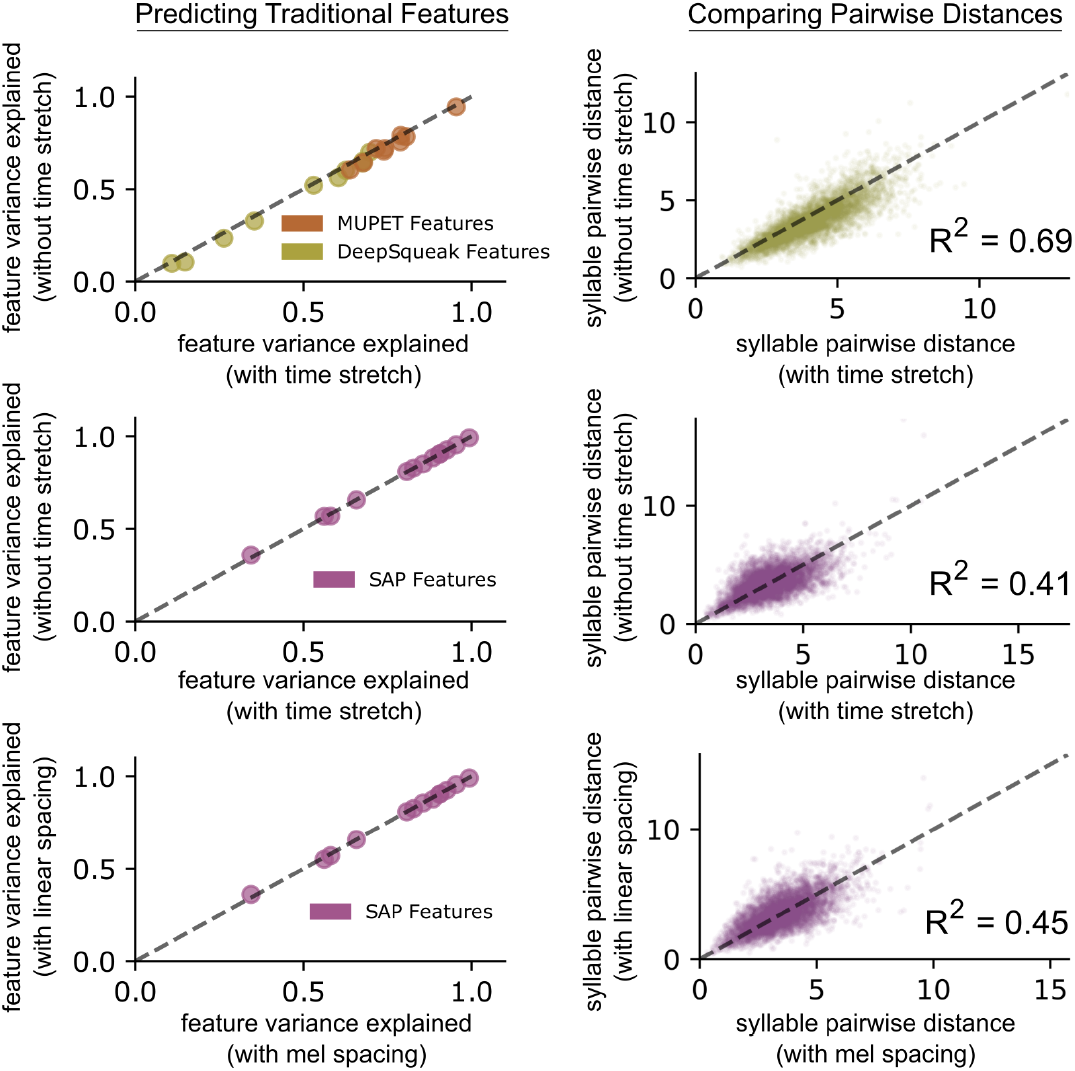
The effect of time stretch and frequency spacing parameters. **top row:** Two VAEs are trained on the mouse USV syllables from Figure 4d, one with time stretching to expand the spectrograms of short syllables (see Methods), and one without. When using the learned latent features from each model to predict the values of acoustic features calculated by MUPET and DeepSqueak, we observe a small but consistent performance gain using time stretching (left column). Additionally, we find a good correspondence between pairwise Euclidean distances in the two learned feature spaces (right column), indicating the two latent spaces have similar geometries. **middle row:** Repeating the same comparison with zebra finch song syllables from Figure 4a-c, we find fairly consistent pairwise distances across the two latent spaces (right) and no substantial effect on the performance of predicting acoustic features calculated by SAP (left). **bottom row:** Two VAEs trained on the same zebra finch song syllables, one with linearly-spaced spectrogram frequency bins and the other with mel-spaced frequency bins, have fairly consistent pairwise distances across the two latent spaces (right) and no substantial effect on the performance of predicting acoustic features calculated by SAP (left).

**Figure S5:**
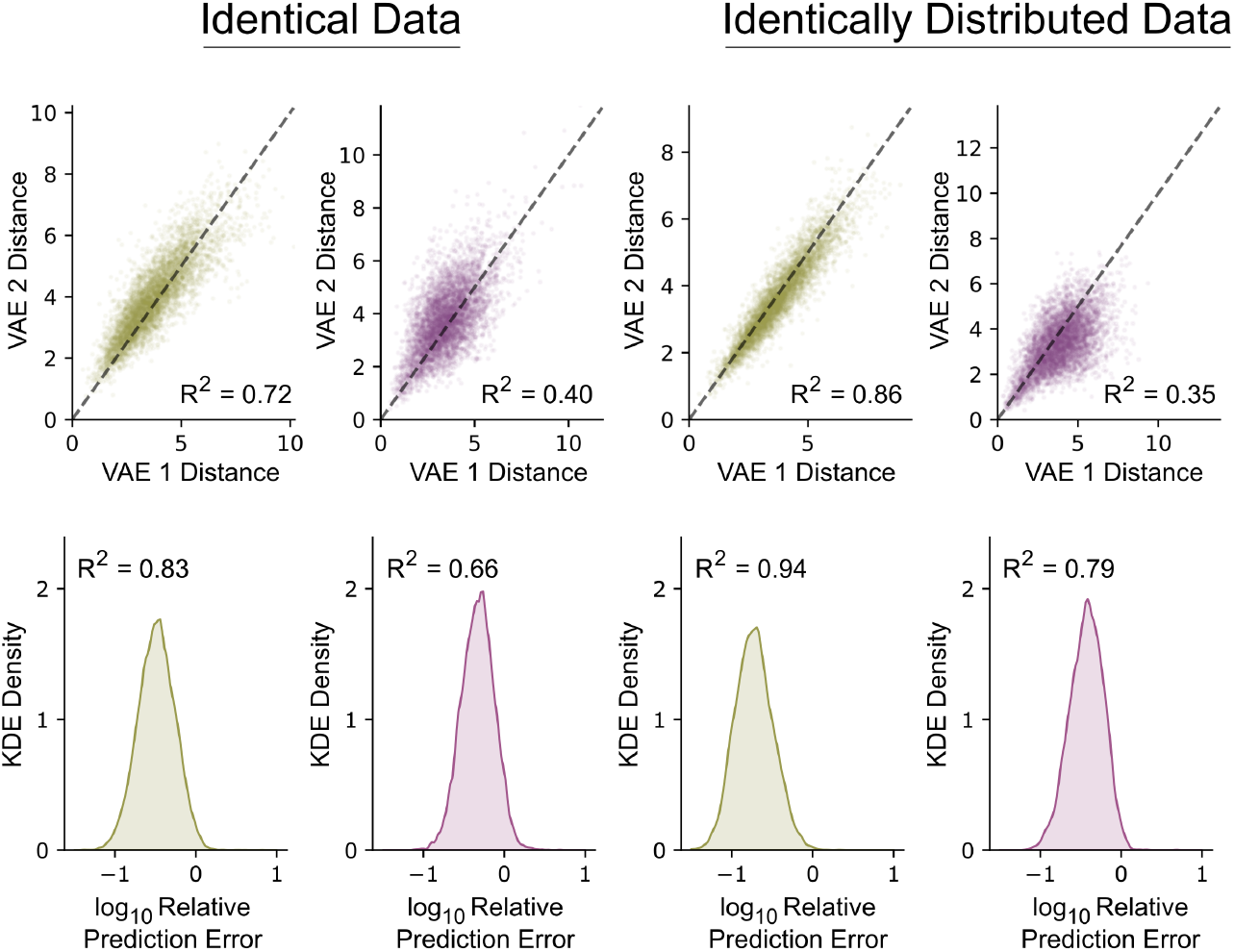
The VAE produces similar latent feature when retraining on identical data and disjoint splits of the same dataset. **top row:** Pairwise latent distances of 5000 random pairs of syllables under separately trained VAEs. **bottom row:** The distribution of errors when predicting the latent means of one VAE from the other using linear regression. Errors are normalized relative to the root-mean-square (RMS) distance from the mean in latent space so that 1 corresponds to 10% of the RMS distance from the mean. For predicting a multivariate *Y* from a multivariate *X*, we report a multivariate *R*^2^ value: 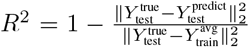. **first column:** Two VAEs are trained on the set of mouse syllables from Figure 2a-c. **second column:** Two VAEs are trained on the set of zebra finch syllables from Figure 4a-c. **third column:** Two VAEs are trained on two disjoint halves of the mouse syllables from Figure 2a-c. **fourth column:** Two VAEs are trained on two disjoint halves of the zebra finch syllables from Figure 4a-c. Note that retraining produces less consistent results for zebra finch syllables, possibly because the relative orientations and positions of well-separated clusters is under-determined by the data.

**Figure S6:**
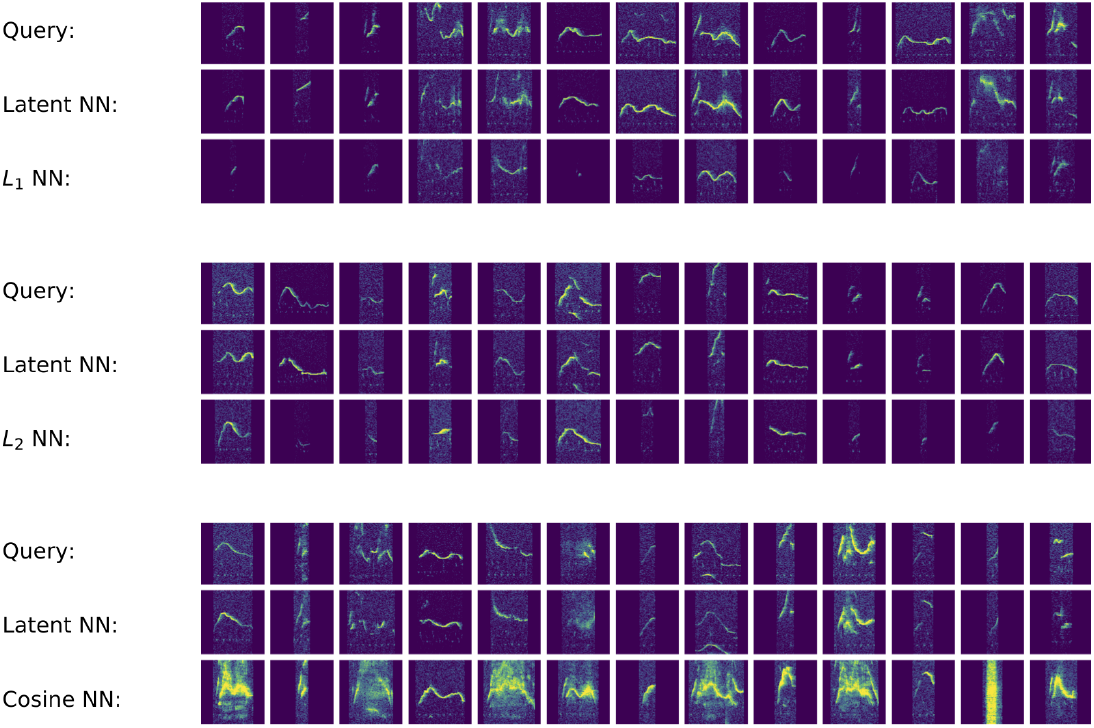
Nearest neighbors returned by distance metrics in spectrogram space exhibit failure modes not found in latent space. **Top block**: Selected query spectrograms, their nearest neighbors in latent space (Euclidean metric), and their nearest neighbors in spectrogram space (*L*_1_ metric). **Middle block**: Same comparison with the *L*_2_ metric in spectrogram space. **Bottom block**: Same comparison with the cosine metric in spectrogram space.

**Figure S7:**
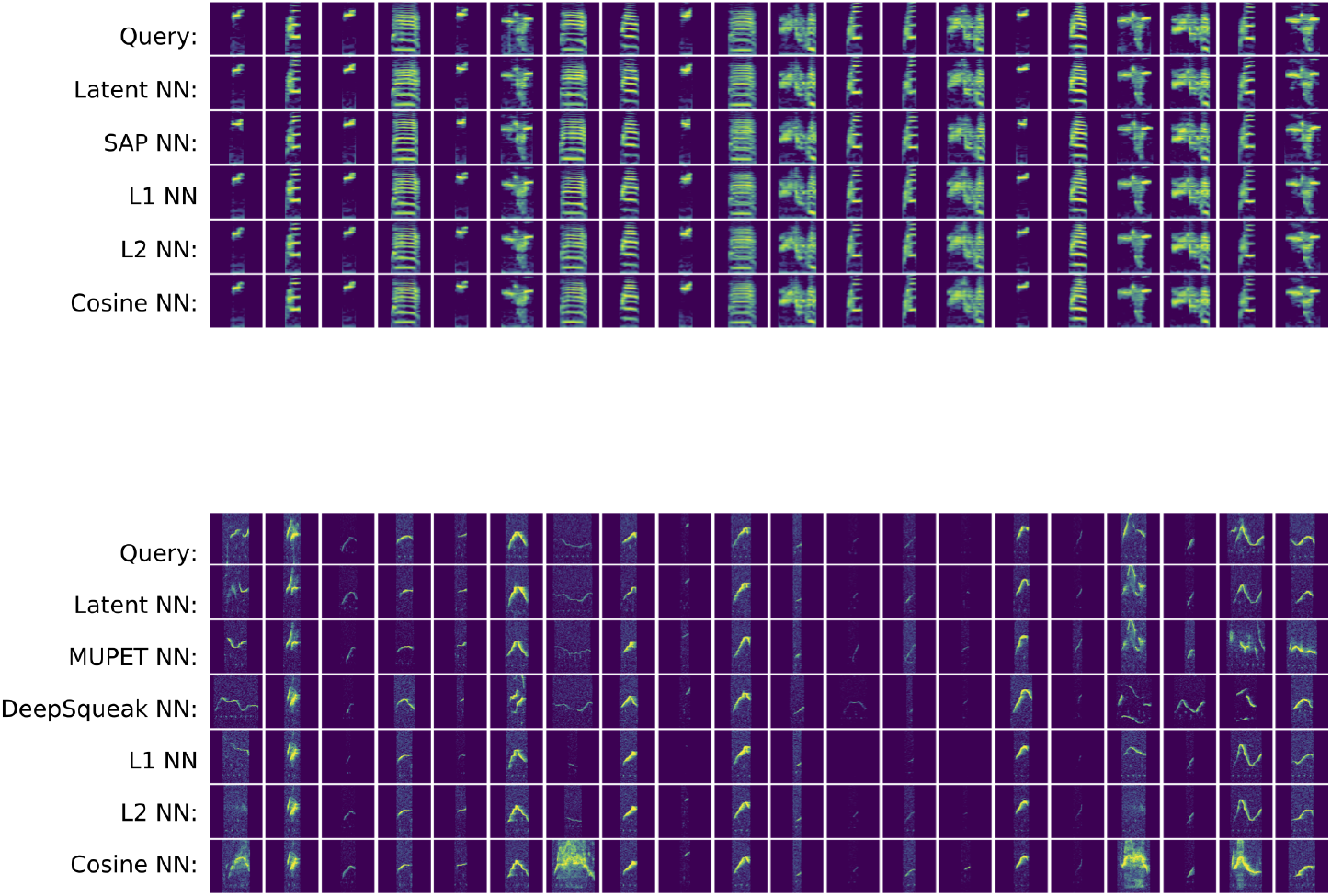
Representative sample of nearest neighbors returned by several feature spaces. **Top block**: Given 20 random zebra finch syllable spectrograms, we find nearest neighbors in five feature spaces: VAE latent space, Sound Analysis Pro feature space, and spectrogram space (Manhattan (L1), Euclidean (L2), and cosine metrics). All methods consistently find nearest neighbors of the same syllable type. **Bottom block**: Given 20 random mouse syllable spectrograms, we find nearest neighbors in six feature spaces: VAE latent space, MUPET feature space, DeepSqueak feature space, and spectrogram space (Manhattan, Euclidean, and cosine metrics). Most methods return mostly similar spectrograms. However, latent features more consistently return good matches than other methods.

**Figure S8:**
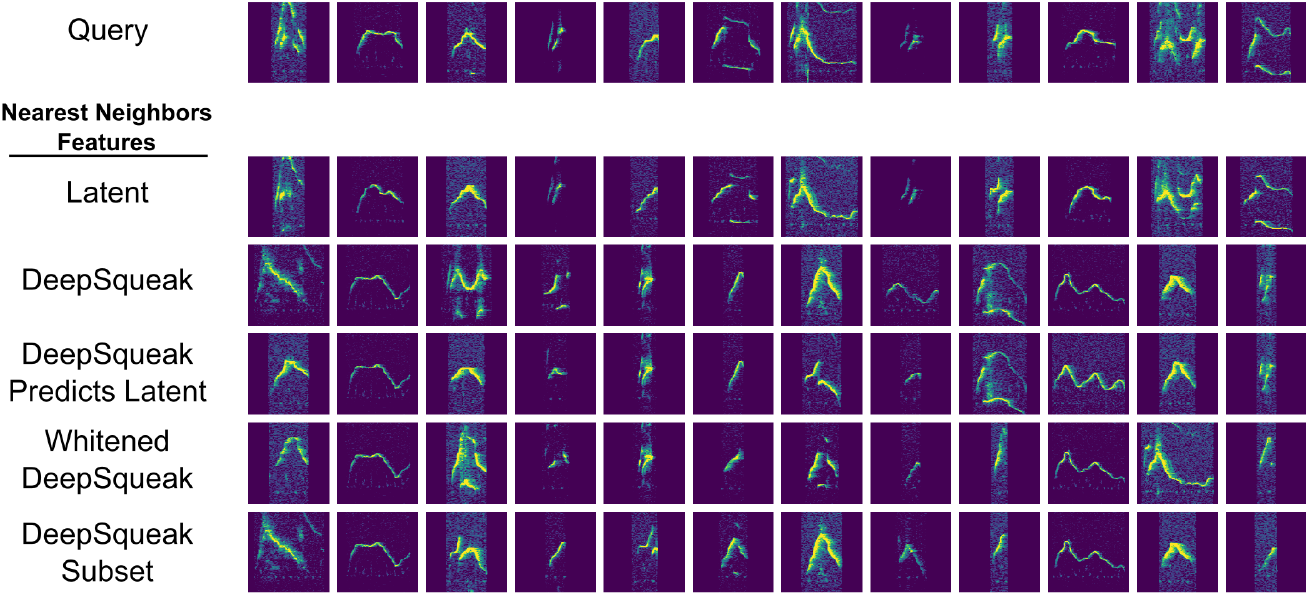
Investigating poor DeepSqueak feature nearest neighbors from Figure 3. **Top row**: Selected query spectrograms from Figure 3. **Lower rows**: Nearest neighbor spectrograms returned by various feature spaces: latent features, DeepSqueak features [7] (standardized, but not whitened), the linear projection of DeepSqueak features that best predicts latent features, whitened DeepSqueak features, and the subset of DeepSqueak features excluding frequency standard deviation, sinuosity, and frequency modulation, the three features most poorly predicted by latent features (Figure S1). All three variants of the DeepSqueak feature set return more visually similar nearest neighbors than the original DeepSqueak feature set for some but not all query spectrograms. In particular, the remaining poor nearest neighbors returned by the linear projection of DeepSqueak features most predictive of latent features suggests DeepSqueak features are insufficient to capture the full acoustic complexity of USV syllables.

**Figure S9:**
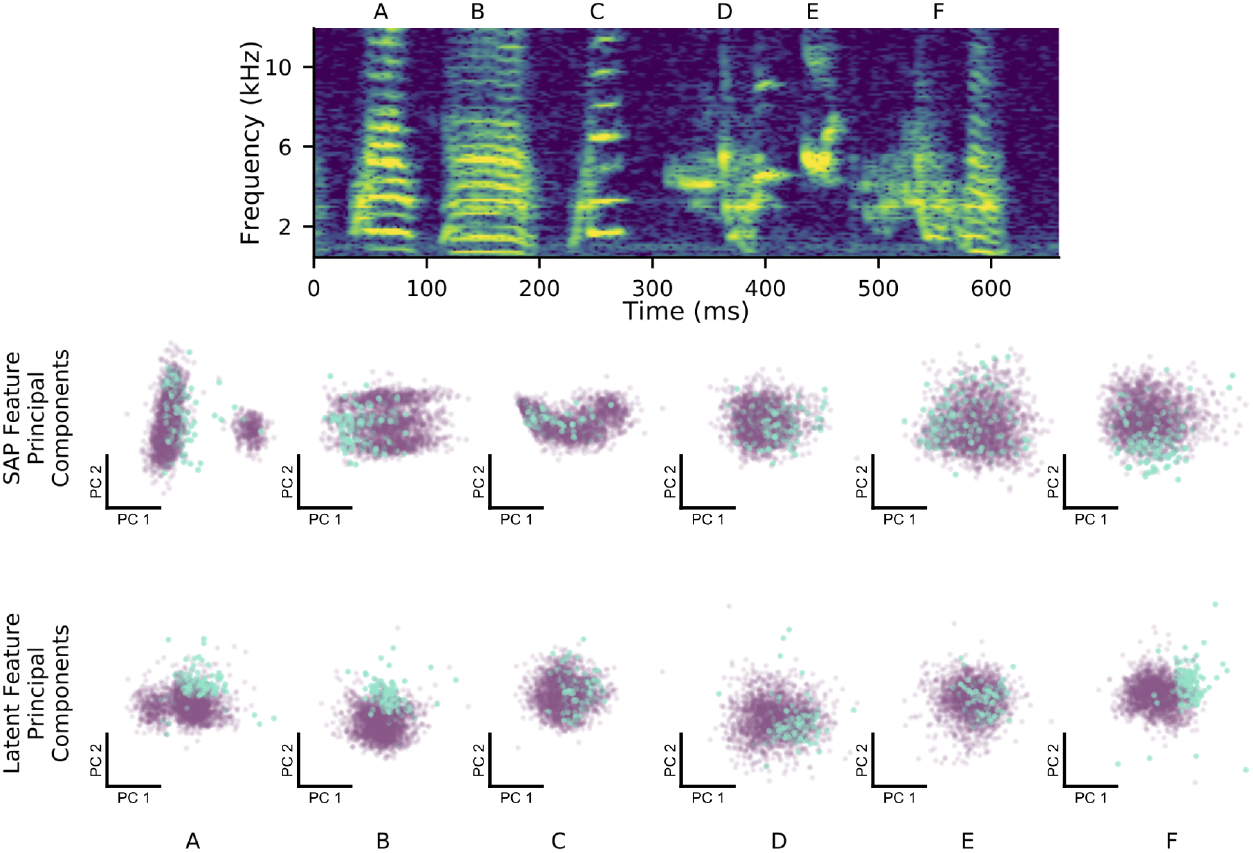
Latent features better represent constricted variability of female-directed zebra finch song. At top is a single rendition of a male zebra finch’s song motif, with individual syllables labeled A-F. The top row of scatterplots shows each syllable over many directed (blue) and undirected (purple) renditions, plotted with respect to the first two principal components of the Sound Analysis Pro acoustic feature space. The bottom row of scatterplots shows the same syllables plotted with respect to the first two principal components of latent feature space. The difference in distributions between the two social contexts is displayed more clearly in the latent feature space, especially for non-harmonic syllables (D,E,F).

**Table S1:** Comparison of feature sets on the downstream task of predicting finch social context (directed vs. undirected context) given acoustic features of single syllables. Classification accuracy, in percent, averaged over 5 disjoint, class-balanced splits of the data is reported. Empirical standard deviation is shown in parentheses. Euclidean distance is used for nearest-neighbor classifiers. Each SAP acoustic feature is independently z-scored as a preprocessing step. Latent feature dimension is truncated when >99% of the feature variance is explained. Random forest (RF) classifiers use 100 trees and the Gini impurity criterion. The multi-layer perceptron (MLP) classifiers are two-layer networks with a hidden layer size of 100, ReLU activations, and an L2 weight regularization parameter “alpha”, trained with ADAM optimization with a learning rate of 10^−3^ for 200 epochs.

| Predicting Finch Social Context (Figure 4a-c) |  |  |  |  |  |
| --- | --- | --- | --- | --- | --- |
|  | Spectrogram |  |  | SAP | Latent |
| | $D = 10$ | $D = 30$ | $D = 100$ | $D = 13$ | $D = 5$ |
| $k$-NN ($k = 3$) | 92.5 (0.2) | 95.3 (0.1) | **97.3 (0.3)** | 93.0 (0.3) | 96.9 (0.2) |
| $k$-NN ($k = 10$) | 93.0 (0.2) | 95.3 (0.2) | 97.1 (0.3) | 93.2 (0.1) | 96.7 (0.2) |
| $k$-NN ($k = 30$) | 92.7 (0.1) | 94.2 (0.2) | 96.0 (0.2) | 92.8 (0.1) | 96.3 (0.1) |
| RF (depth= 10) | 92.6 (0.1) | 92.7 (0.1) | 93.1 (0.1) | 92.8 (0.1) | 94.9 (0.2) |
| RF (depth= 15) | 92.7 (0.1) | 93.2 (0.2) | 93.6 (0.1) | 93.6 (0.2) | 96.1 (0.1) |
| RF (depth= 20) | 92.8 (0.1) | 93.4 (0.1) | 93.8 (0.1) | 93.8 (0.2) | 96.4 (0.2) |
| MLP ($\alpha = 0.1$) | 92.8 (0.4) | 95.4 (0.4) | **97.6 (0.3)** | 92.9 (0.1) | 95.7 (0.1) |
| MLP ($\alpha = 0.01$) | 92.9 (0.3) | 95.4 (0.3) | **97.5 (0.2)** | 93.1 (0.2) | 96.2 (0.1) |
| MLP ($\alpha = 0.001$) | 92.7 (0.6) | 95.2 (0.5) | **97.5 (0.2)** | 93.0 (0.2) | 96.3 (0.0) |

**Table S2:**
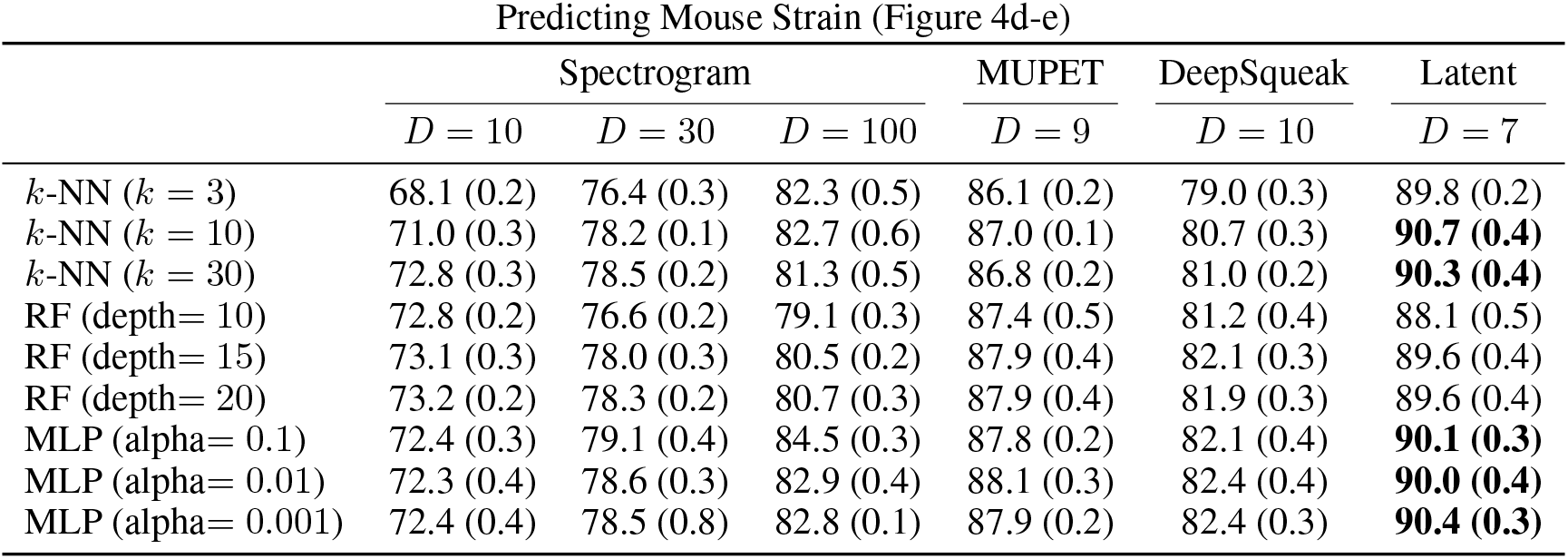
Comparison of feature sets on the downstream task of predicting mouse strain (C57 vs. DBA) given acoustic features of single syllables. Classification accuracy, in percent, averaged over 5 disjoint, class-balanced splits of the data is reported. Empirical standard deviation is shown in parentheses. Euclidean distance is used for nearest-neighbor classifiers. Each MUPET and DeepSqueak acoustic feature is independently z-scored as a preprocessing step. Latent features dimension is truncated when >99% of the feature variance is explained. Random forest (RF) classifiers use 100 trees and the Gini impurity criterion. The multi-layer perceptron (MLP) classifiers are two-layer networks with a hidden layer size of 100, ReLU activations, and an L2 weight regularization parameter “alpha”, trained with ADAM optimization with a learning rate of 10^−3^ for 200 epochs.

**Table S3:** Comparison of feature sets on the downstream task of predicting mouse identity given acoustic features of single syllables. Classification accuracy, in percent, averaged over 5 disjoint, class-balanced splits of the data is reported. A class-weighted log-likelihood loss is targeted to help correct for class imbalance. Empirical standard deviation is shown in parentheses. Each MUPET acoustic feature is independently z-scored as a preprocessing step. Latent feature principal components are truncated when >99% of the feature variance is explained. Multi-layer perceptron (MLP) classifiers are two-layer networks with a hidden layer size of 100, ReLU activations, and an L2 weight regularization parameter “alpha”, trained with ADAM optimization with a learning rate of 10^−^3 for 200 epochs. Chance performance is 2.8% for top-1 accuracy and 13.9% for top-5 accuracy.

| Predicting Mouse Identity (Figure 4f) |  |  |  |  |  |
| --- | --- | --- | --- | --- | --- |
| top-1 accuracy | Spectrogram |  |  | MUPET | Latent |
| | $D = 10$ | $D = 30$ | $D = 100$ | $D = 9$ | $D = 8$ |
| MLP (alpha= 0.01) | 9.9 (0.2) | 14.9 (0.2) | 20.4 (0.4) | 14.7 (0.2) | 17.0 (0.3) |
| MLP (alpha= 0.001) | 10.8 (0.1) | 17.3 (0.4) | **25.3 (0.3)** | 19.0 (0.3) | 22.7 (0.5) |
| MLP (alpha= 0.0001) | 10.7 (0.2) | 17.3 (0.3) | **25.1 (0.3)** | 20.6 (0.4) | 24.0 (0.2) |
| top-5 accuracy |  |  |  |  |  |
| MLP (alpha= 0.01) | 36.6 (0.4) | 45.1 (0.5) | 55.0 (0.3) | 46.5 (0.3) | 49.9 (0.4) |
| MLP (alpha= 0.001) | 38.6 (0.2) | 50.7 (0.6) | **62.9 (0.4)** | 54.0 (0.2) | 59.2 (0.6) |
| MLP (alpha= 0.0001) | 38.7 (0.5) | 50.8 (0.3) | **63.2 (0.4)** | 57.3 (0.4) | 61.6 (0.4) |

**Figure S10:**
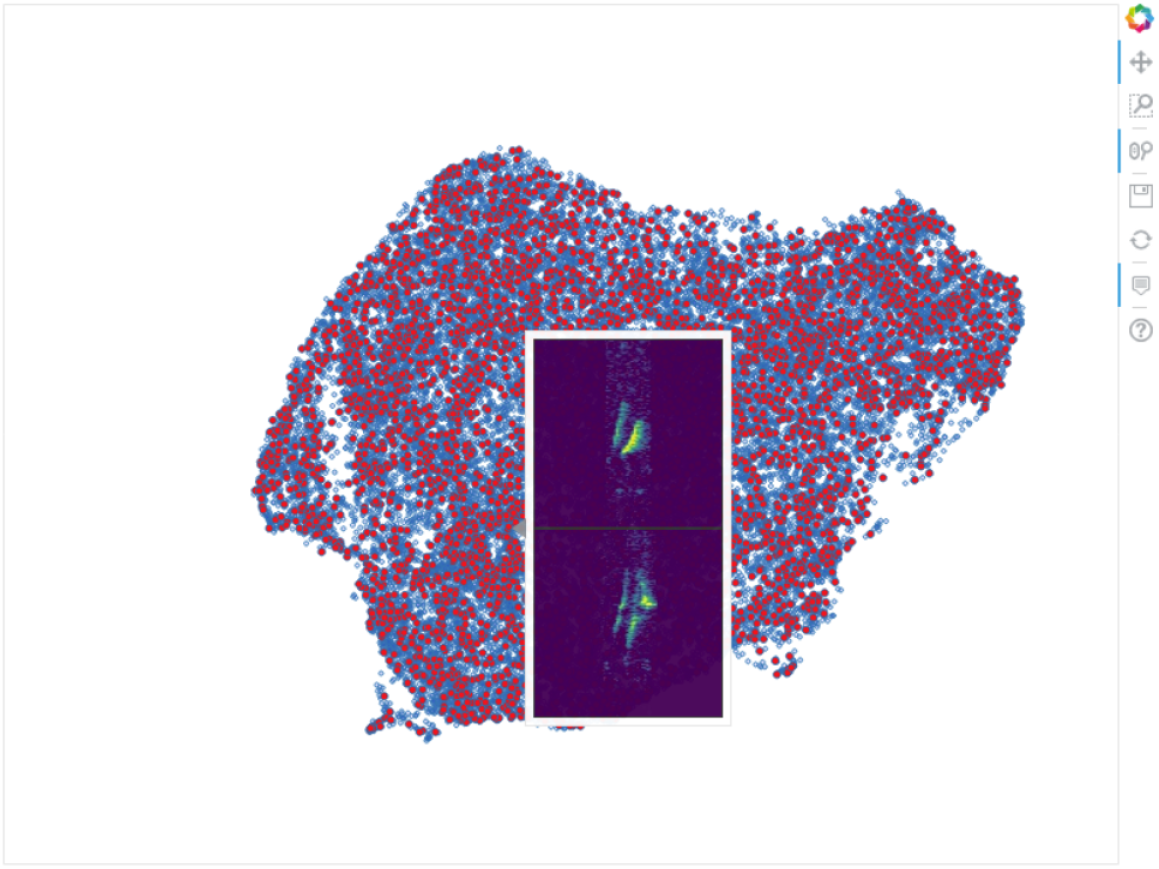
“Atlas” of mouse USVs. This screenshot shows an interactive version of Figure 4d in which example spectrograms are displayed as tooltips when a cursor hovers over the plot. A version of this plot is hosted at the following web address: https://pearsonlab.github.io/research.html#mouse_tooltip

**Figure S11:**
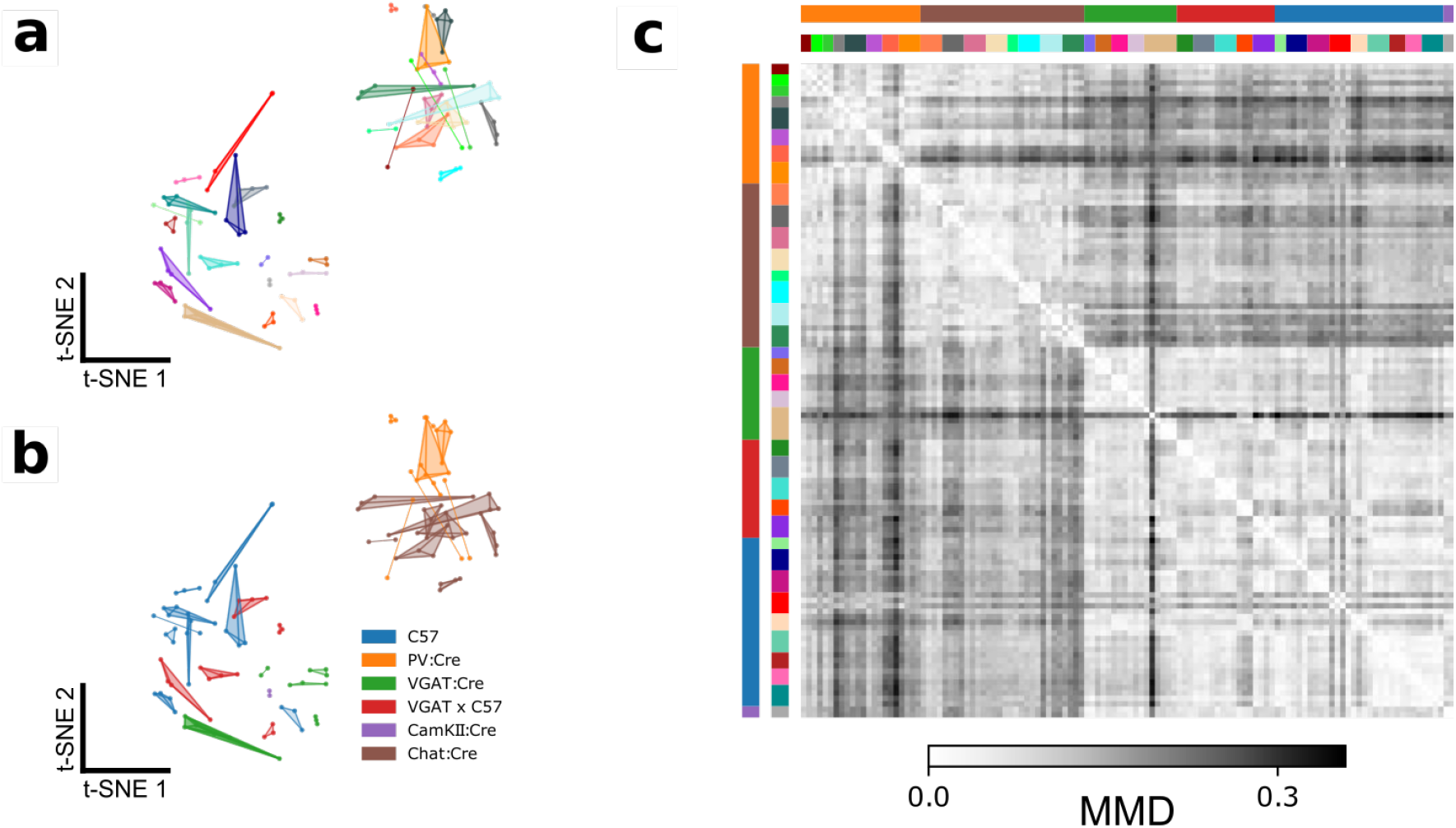
Details of Figure 4f. **a)** t-SNE representation of similarities between syllable repertoires, where distance metric is estimated MMD between latent syllable distributions (reproduction of Figure 4f). Each scatter represents the USV syllable repertoire of a single recording session. Recordings of the same mice across different days are connected and colored identically. Distances between points represent the similarity in vocal repertoires, with closer points more similar. Note that the most mice have similar repertoires across days, indicated by the close proximity of connected scatterpoints. **b)** The same plot as **a**, colored by the genetic background of each mouse. Note that the two primary clusters of USV syllable repertoires correspond to two distinct sets of genetic backgrounds. **c)** The full pairwise MMD matrix between USV repertoires from individual recording sessions. The dataset contains 36 individuals, 118 recording sessions, and 156,180 total syllables. The two main clusters separating the PV:Cre and Chat:Cre mice from the other backgrounds is apparent as the large two-by-two checkerboard pattern. Colors at the top and left sides indicate individual and genetic background information, with colors matching those in panels **a** and **b**.

**Figure S12:**
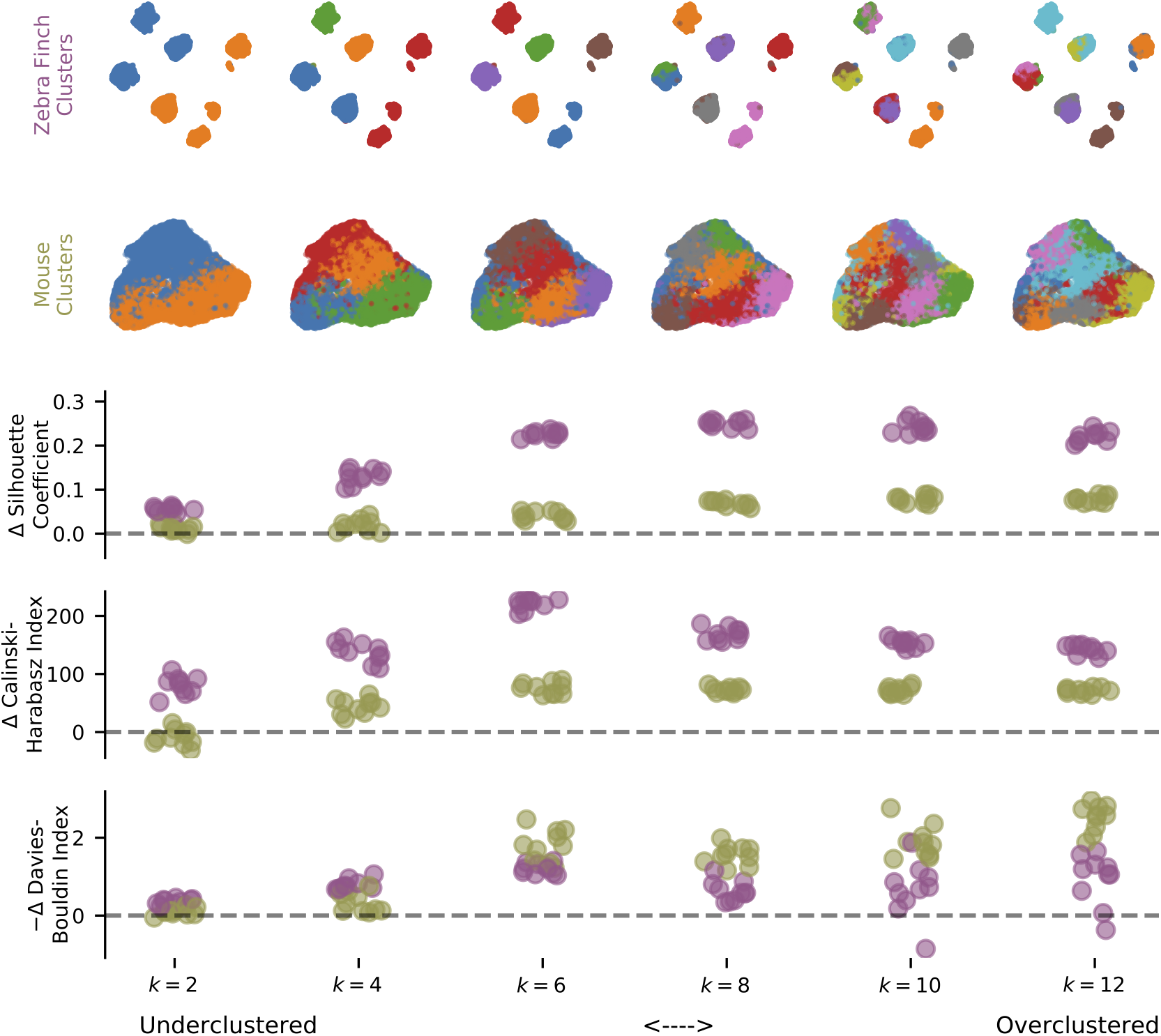
Three unsupervised clustering metrics evaluated on the latent description of zebra finch song syllables (Figure 5a) and mouse USV syllables (Figure 5b) as the number of components, *k*, varies from 2 to 12. Clustering metrics are reported relative to moment-matched Gaussian noise (see Methods) with a possible sign change so that higher scores indicate more clustering.

**Figure S13:**
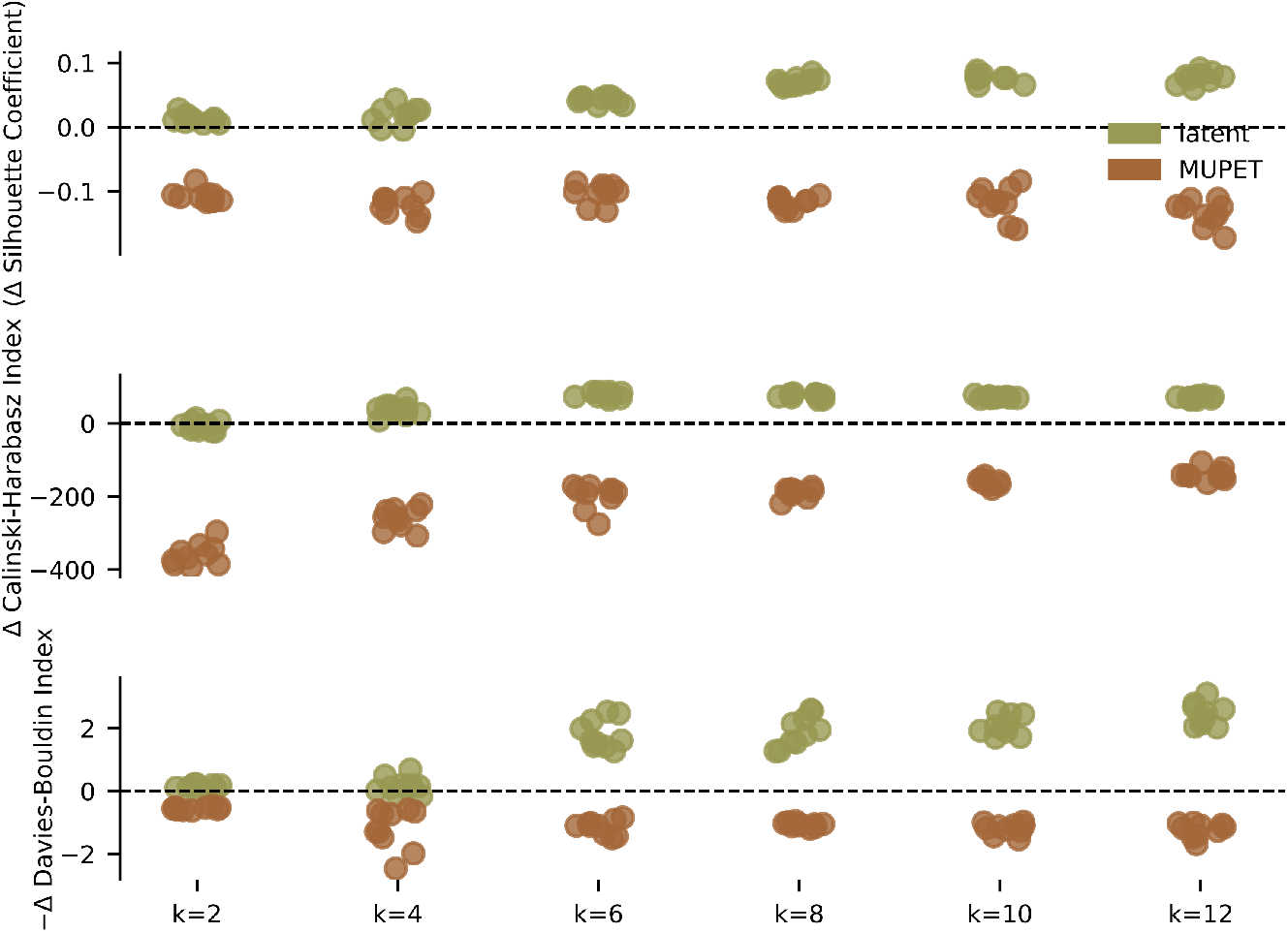
Three unsupervised clustering metrics evaluated on latent and MUPET features of mouse USV syllables (Figure 5b) as the number of components, *k*, varies from 2 to 12. Clustering metrics are reported relative to moment-matched Gaussian noise (see Methods) with a possible sign change so that higher scores indicate more clustering. Latent features are consistently judged to produce better clustering than MUPET feature. Additionally, MUPET features are consistently judged to be less clustered than moment-matched Gaussian noise. Compare to Figure S14.

**Figure S14:**
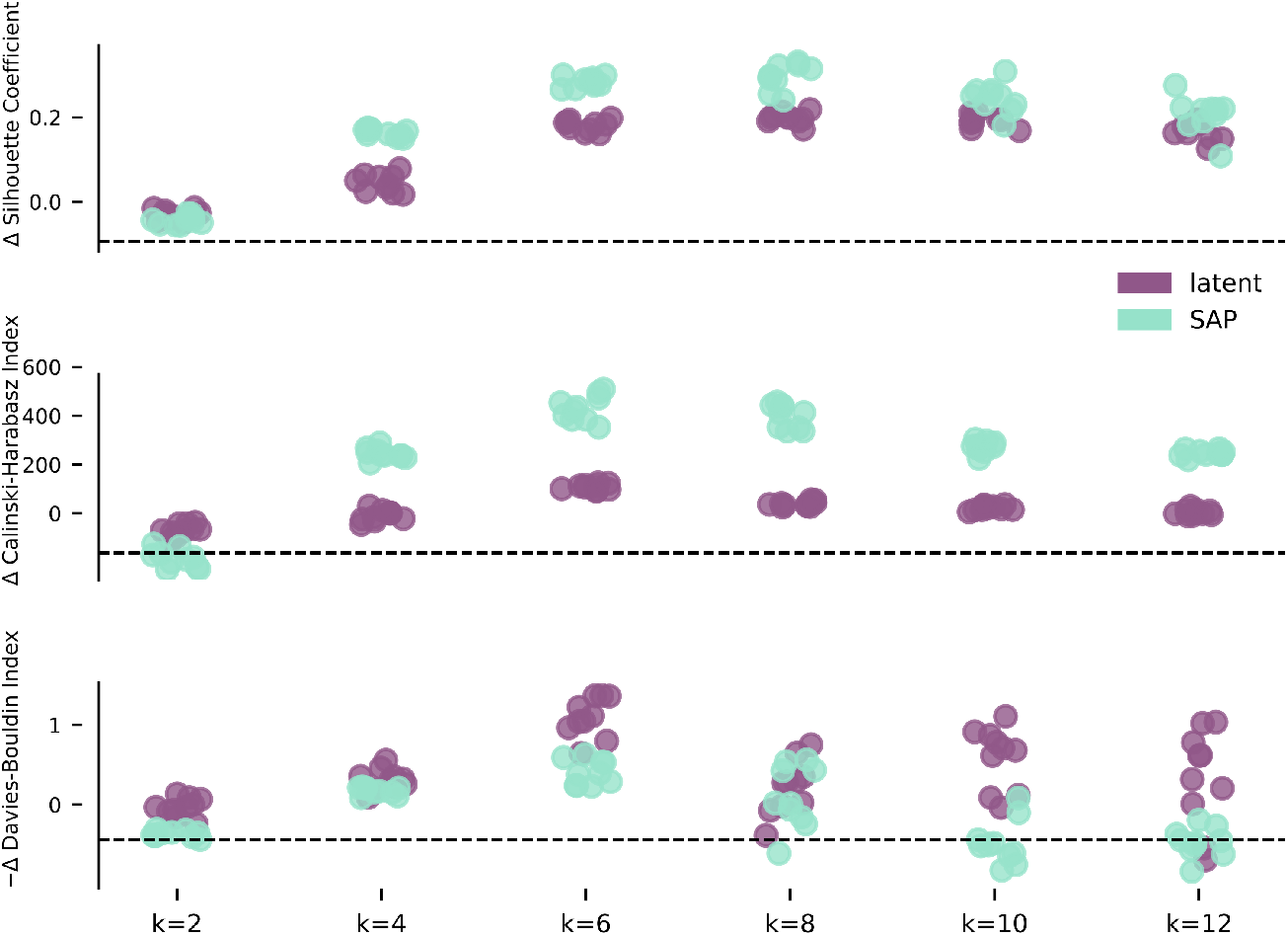
Three unsupervised clustering metrics evaluated on latent and SAP features of zebra finch song syllables (Figure 5a) as the number of components, *k*, varies from 2 to 12. Clustering metrics are reported relative to moment-matched Gaussian noise (see Methods) with a possible sign change so that higher scores indicate more clustering. Both feature sets admit clusters that are consistently judged to be more clustered than moment-matched Gaussian noise.

**Figure S15:**
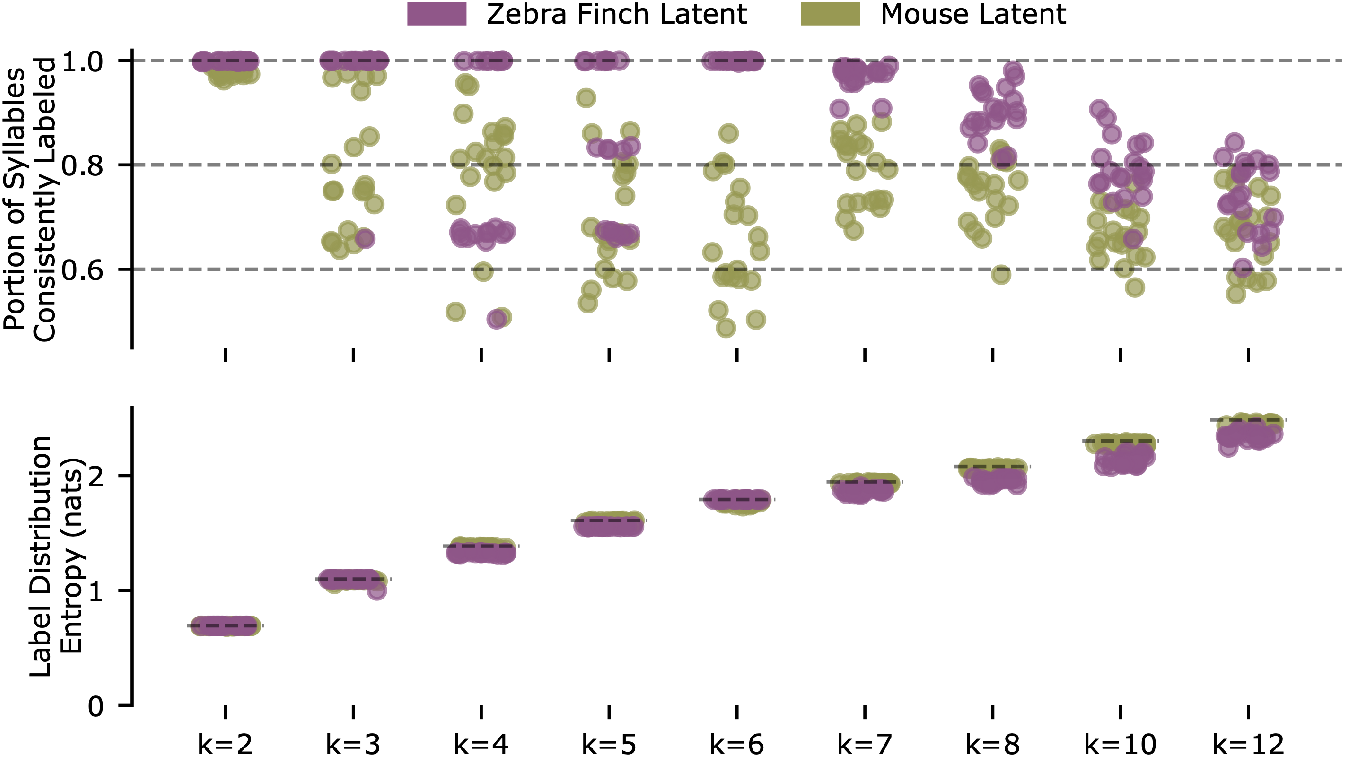
Repeated clustering with Gaussian Mixture Models (GMMs) produces reliable clusters for zebra finch syllable latent features with six clusters, but not for mouse syllable latent features with more than two clusters. Both sets of syllable latent descriptions (zebra finch, 14270 syllables; mouse, 15712 syllables) are repeatedly split into thirds. The first and second splits are used to train Gaussian Mixture Models (GMMs, full covariance, best of 5 fits, fit via expectation maximization), which are used to predict labels on the third split. Given these predicted labels and a matching of labels between the two GMMs, a syllable can be considered consistently labeled if it is assigned the same label class by the two GMMs. The Hungarian method is used to find the label matching that maximizes the portion of consistently labeled syllables. **top row)** The portion of consistently labeled syllables for 20 repetitions of this procedure is shown for varying numbers of clusters, *k*. Note that zebra finch syllables achieve near-perfect consistency for six clusters, the number of clusters found by hand labeling (syllables A-F in Figure 6c), and this consistency degrades with more clusters. In contrast, the consistency of mouse USV clusters is poor for *k* > 2. Somewhat surprisingly, the clustering is very consistent for two clusters (*k* = 2). To test whether this is a trivial effect of rarely used clusters, we calculated the entropy of the empirical label distributions (**bottom row**). We find in each case, and specifically the *k* = 2 case, that the empirical distribution entropy is close to the maximum possible entropy (plotted as dashed horizontal lines), indicating that all clusters are frequently used. While consistent with mouse USVs forming two clusters, this result is not sufficient proof of syllable clustering or even bimodality. Yet, for cluster identity to be a practical syllable descriptor, clusters should be readily identifiable from the data. Thus the combination of VAE-identified latent features and Gaussian clusters does not appear to be a suitable description of mouse USVs for *k >* 2 clusters. Note that the portion of consistently labeled zebra finch syllables for *k* < 6 clusters form well-defined bands at multiples of 1, indicating that cluster structure is so well defined by the data that the GMMs do not split individual clusters across multiple Gaussian components.

**Figure S16:**
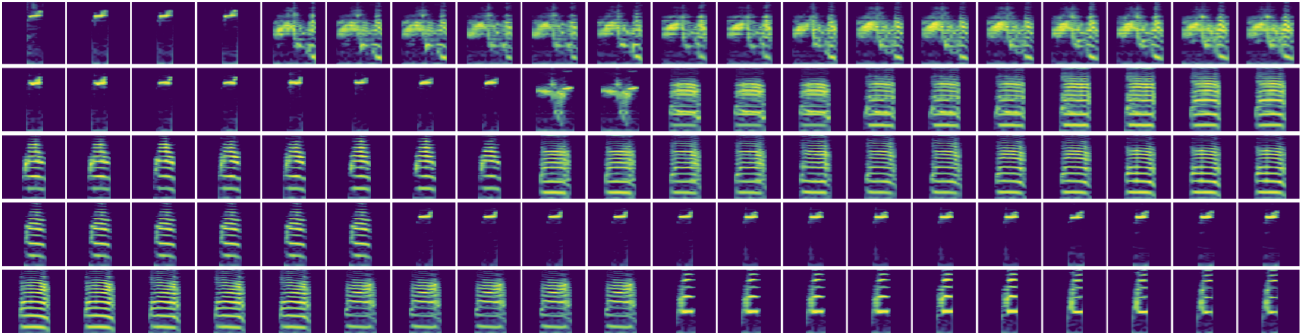
Absence of continuous interpolations between zebra finch song syllables. Each row displays two random zebra finch syllables of different syllable types at either end and an attempted smooth interpolation between the two. Interpolating spectrograms are those with the closest latent features along a linear interpolation in latent space. Note the discontinuous jump in each attempted interpolation, which is expected given that adult zebra finch syllables are believed to be well-clustered. Compare with Figure 5e, which shows continuous variation in mouse USVs.

**Figure S17:**
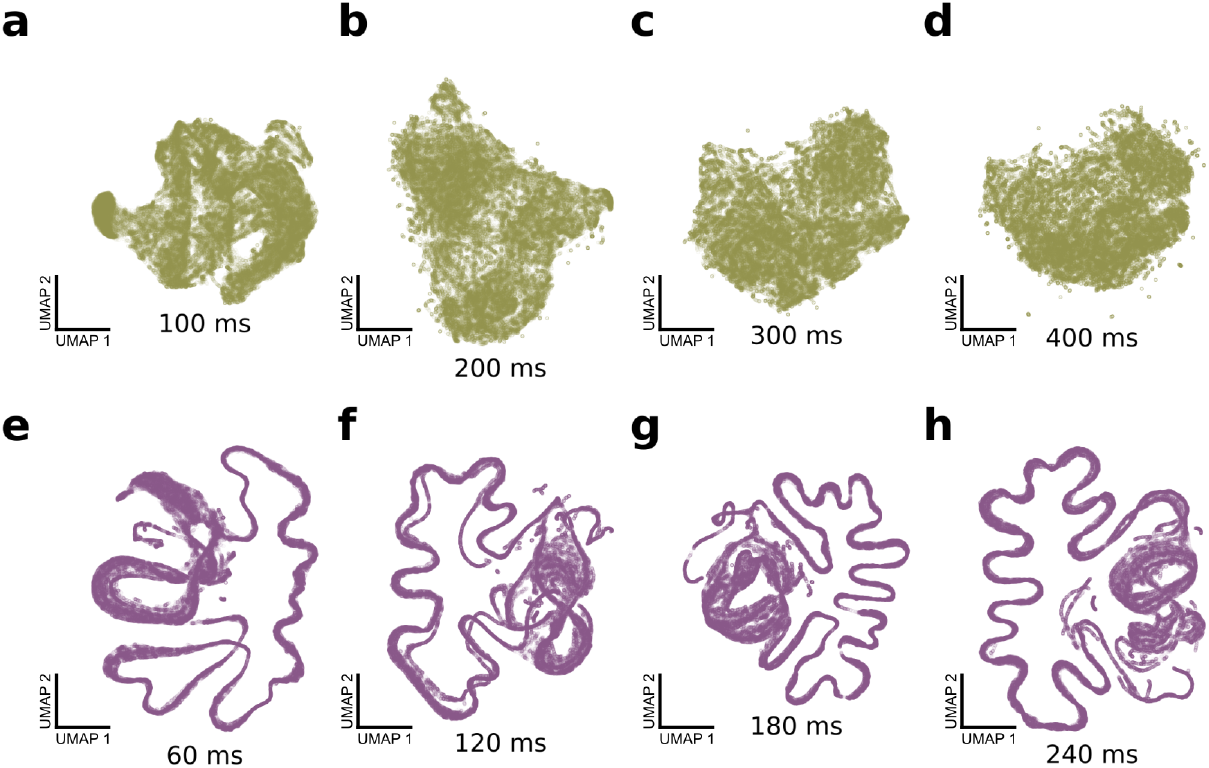
Qualitatively similar shotgun VAE latent projections are achieved with a wide range of window durations. **a-d**) UMAP projections of 100,000 windows of mouse USVs, with window durations of 100, 200, 300, and 400ms. Compare with Figure 6a. **e-h**) UMAP projections of 100,000 windows of zebra finch song motifs, with window durations of 60, 120, 180, and 240ms. Compare with Figure 6b.

**Figure S18:**
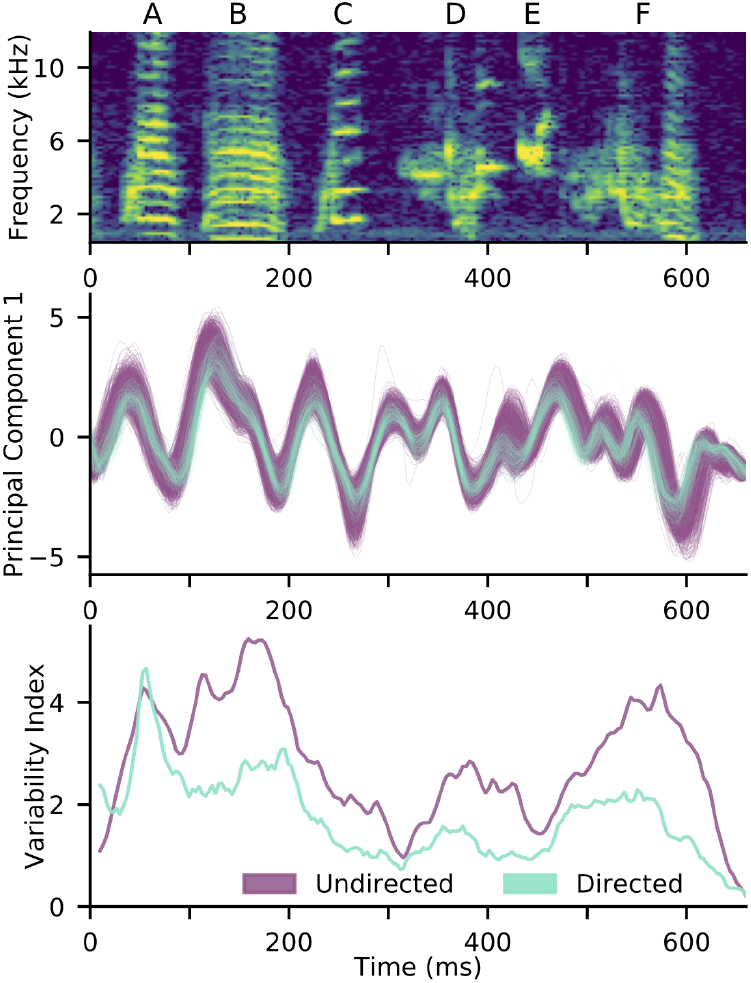
Non-timewarped version of Figure 6c-e. As in Figure 6d, there is reduced variability in the first latent principal component for directed song compared to undirected song. The overall variability reduction is quantified by the variability index (see Methods), reproducing the reduced variability of directed song found in Figure 6e. Note that the directed traces lag the undirected traces at the beginning of the motif and lead at the end of the motif due to their faster tempo, which is uncorrected in this version of the analysis.

**Figure S19:**
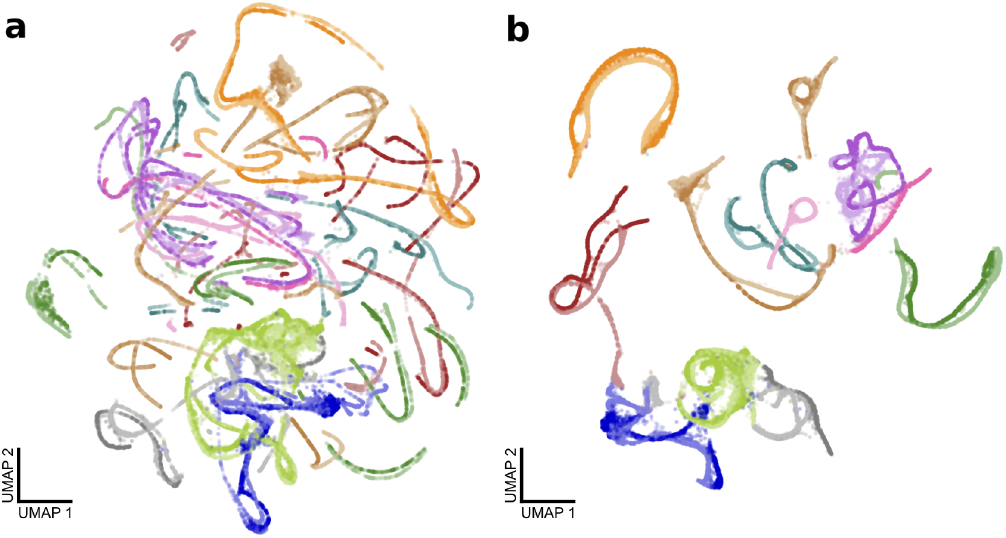
The effect of the modified UMAP distance matrix from Figure 7d-e. **a)** A UMAP projection of shotgun VAE latent means with a standard Euclidean metric. Colors represent birds as in Figure 7. **b)** A UMAP projection of the same latent means with a modified metric to discourage strands from the same motif rendition from splitting (see Methods). This is a reproduction of Figure 7d.

**Figure S20:**
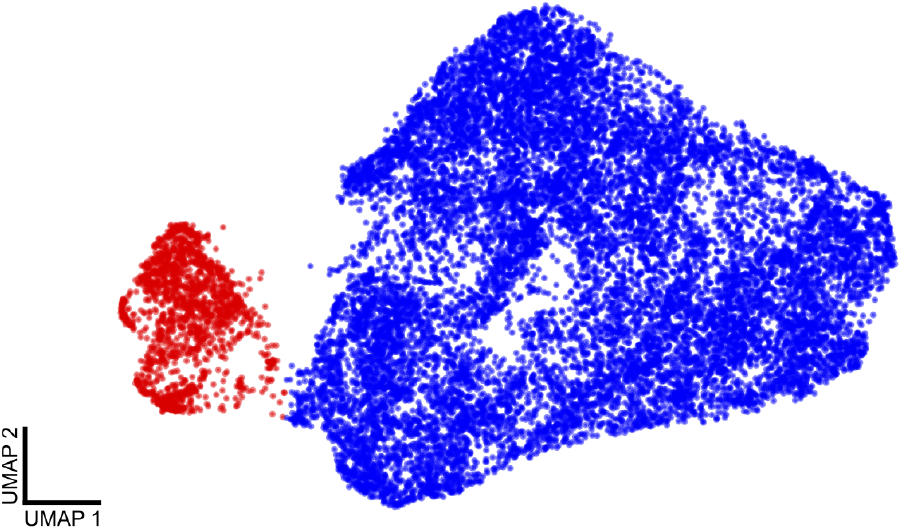
Removing noise from single mouse recordings (see Recordings). Above is a UMAP projection of all detected USV syllables. The false positives (red) cluster fairly well, so they were removed from further analysis. Of the 17,400 total syllables detected, 15,712 (blue) remained after removing the noise cluster.

**Figure S21:**
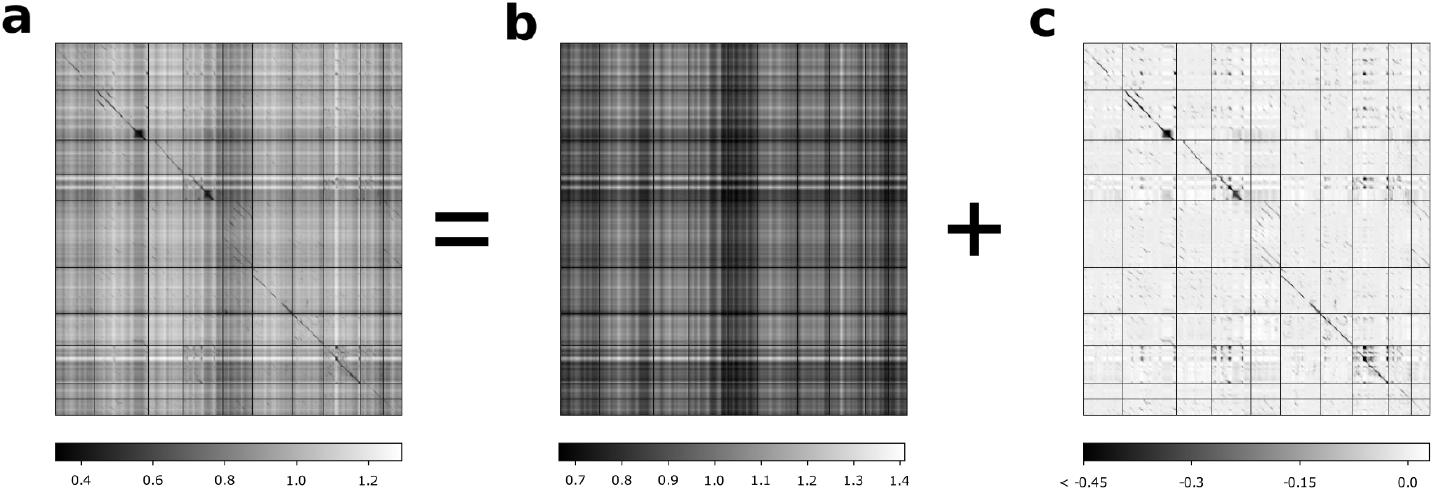
For the shotgun VAE pupil/tutor analysis presented in Figure 7, an MMD matrix (**a**) is decomposed into a rank-one component (**b**) and a residual component (**c**). The residual component is shown in Figure 7f for visual clarity. The decomposition is performed by MAP estimation assuming a flat prior on the rank-1 matrix and independent Laplace errors on the residual component.

**Figure S22:**
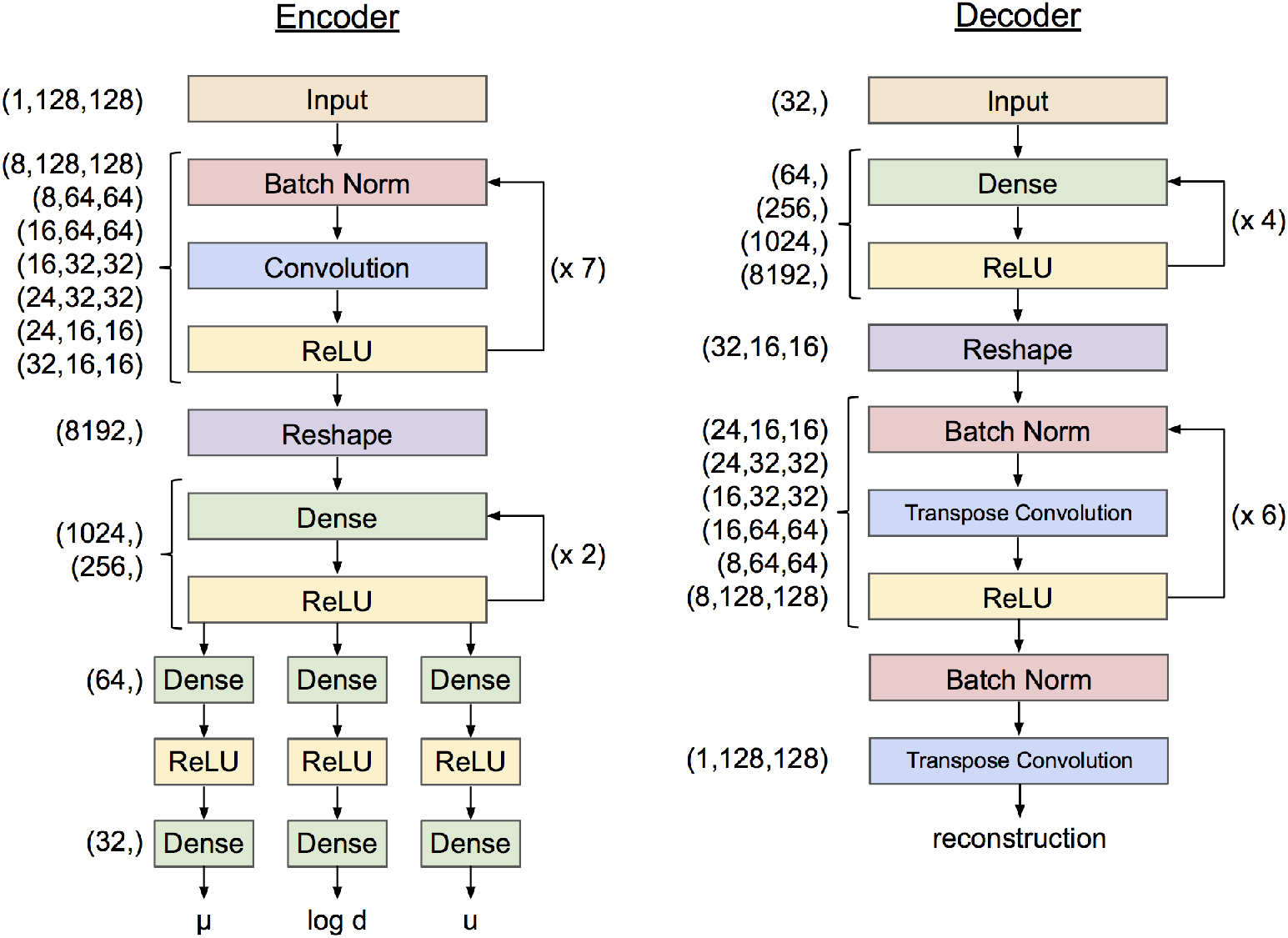
VAE network architecture. The architecture outlined above was used for all training runs. The looping arrows at the right of the encoder and decoder denote repeated sequences of layer types, not recurrent connections. For training details see Methods. For implementation details, see: https://github.com/pearsonlab/autoencoded-vocal-analysis

## References

[1] Mupet wiki. https://github.com/mvansegbroeck/mupet/wiki/MUPET-wiki. Accessed: 2019-09-07.

[2] Julia C Berryman. Guinea-pig vocalizations: their structure, causation and function. Zeitschrift für Tierpsychologie, 41(1):80–106, 1976.

[3] Christopher P Burgess, Irina Higgins, Arka Pal, Loic Matthey, Nick Watters, Guillaume Desjardins, and Alexander Lerchner. Understanding disentangling in *β*-vae. arXiv preprint arXiv:1804.03599, 2018.

[4] Zachary D Burkett, Nancy F Day, Olga Peñagarikano, Daniel H Geschwind, and Stephanie A White. Voice: A semi-automated pipeline for standardizing vocal analysis across models. Scientific reports, 5:10237, 2015.

[5] Tadeusz Caliński and Jerzy Harabasz. A dendrite method for cluster analysis. Communications in Statistics-theory and Methods, 3(1):1–27, 1974.

[6] Jonathan Chabout, Abhra Sarkar, David B Dunson, and Erich D Jarvis. Male mice song syntax depends on social contexts and influences female preferences. Frontiers in behavioral neuroscience, 9:76, 2015.

[7] Kevin R Coffey, Russell G Marx, and John F Neumaier. Deepsqueak: a deep learning-based system for detection and analysis of ultrasonic vocalizations. Neuropsychopharmacology, 44(5):859, 2019.

[8] Bin Dai, Yu Wang, John Aston, Gang Hua, and David Wipf. Connections with robust pca and the role of emergent sparsity in variational autoencoder models. The Journal of Machine Learning Research, 19(1):1573–1614, 2018.

[9] David L Davies and Donald W Bouldin. A cluster separation measure. IEEE transactions on pattern analysis and machine intelligence, (2):224–227, 1979.

[10] Sébastien Derégnaucourt, Partha P Mitra, Olga Fehér, Carolyn Pytte, and Ofer Tchernichovski. How sleep affects the developmental learning of bird song. Nature, 433(7027):710, 2005.

[11] Michale S Fee and Jesse H Goldberg. A hypothesis for basal ganglia-dependent reinforcement learning in the songbird. Neuroscience, 198:152–170, 2011.

[12] Olga Fehér, Haibin Wang, Sigal Saar, Partha P Mitra, and Ofer Tchernichovski. De novo establishment of wild-type song culture in the zebra finch. Nature, 459(7246):564, 2009.

[13] Simone Gaub, Matthias Groszer, Simon E Fisher, and Günter Ehret. The structure of innate vocalizations in foxp2-deficient mouse pups. Genes, Brain and Behavior, 9(4):390–401, 2010.

[14] Arthur Gretton, Karsten M Borgwardt, Malte J Rasch, Bernhard Schölkopf, and Alexander Smola. A kernel two-sample test. Journal of Machine Learning Research, 13(Mar):723–773, 2012.

[15] Kurt Hammerschmidt, Konstantin Radyushkin, Hannelore Ehrenreich, and Julia Fischer. The structure and usage of female and male mouse ultrasonic vocalizations reveal only minor differences. PloS one, 7(7):e41133, 2012.

[16] Stav Hertz, Benjamin Weiner, Nisim Perets, and Michael London. Temporal structure of mouse courtship vocalizations facilitates syllable labeling. Communications Biology, 3(1):1–13, 2020.

[17] Irina Higgins, Loic Matthey, Arka Pal, Christopher Burgess, Xavier Glorot, Matthew Botvinick, Shakir Mohamed, and Alexander Lerchner. beta-vae: Learning basic visual concepts with a constrained variational framework. ICLR, 2(5):6, 2017.

[18] Timothy E Holy and Zhongsheng Guo. Ultrasonic songs of male mice. PLoS biology, 3(12):e386, 2005.

[19] A Ivanenko, Paul Watkins, MAJ van Gerven, K Hammerschmidt, and B Englitz. Classifying sex and strain from mouse ultrasonic vocalizations using deep learning. PLoS computational biology, 16(6):e1007918, 2020.

[20] Anil K Jain, M Narasimha Murty, and Patrick J Flynn. Data clustering: a review. ACM computing surveys (CSUR), 31(3):264–323, 1999.

[21] Mimi H Kao and Michael S Brainard. Lesions of an avian basal ganglia circuit prevent context-dependent changes to song variability. Journal of neurophysiology, 96(3):1441–1455, 2006.

[22] Arik Kershenbaum, Daniel T Blumstein, Marie A Roch, Çağlar Akçay, Gregory Backus, Mark A Bee, Kirsten Bohn, Yan Cao, Gerald Carter, Cristiane Cäsar, et al. Acoustic sequences in non-human animals: a tutorial review and prospectus. Biological Reviews, 91(1):13–52, 2016.

[23] Ilyes Khemakhem, Diederik P Kingma, and Aapo Hyvärinen. Variational autoencoders and nonlinear ica: A unifying framework. arXiv preprint arXiv:1907.04809, 2019.

[24] Diederik P Kingma and Jimmy Ba. Adam: A method for stochastic optimization. arXiv preprint arXiv:1412.6980, 2014.

[25] Diederik P Kingma and Max Welling. Auto-encoding variational bayes. arXiv preprint arXiv:1312.6114, 2013.

[26] Sepp Kollmorgen, Richard HR Hahnloser, and Valerio Mante. Nearest neighbours reveal fast and slow components of motor learning. Nature, 577(7791):526–530, 2020.

[27] Zhifeng Kong, Wei Ping, Jiaji Huang, Kexin Zhao, and Bryan Catanzaro. Diffwave: A versatile diffusion model for audio synthesis. arXiv preprint arXiv:2009.09761, 2020.

[28] Christos Louizos, Kevin Swersky, Yujia Li, Max Welling, and Richard Zemel. The variational fair autoencoder. arXiv preprint arXiv:1511.00830, 2015.

[29] Laurens van der Maaten and Geoffrey Hinton. Visualizing data using t-sne. Journal of machine learning research, 9(Nov):2579–2605, 2008.

[30] Yael Mandelblat-Cerf and Michale S Fee. An automated procedure for evaluating song imitation. PloS one, 9(5):e96484, 2014.

[31] Leland McInnes, John Healy, and James Melville. Umap: Uniform manifold approximation and projection for dimension reduction. arXiv preprint arXiv:1802.03426, 2018.

[32] David G Mets and Michael S Brainard. An automated approach to the quantitation of vocalizations and vocal learning in the songbird. PLoS computational biology, 14(8):e1006437, 2018.

[33] Jacqueline R Miller and Mark D Engstrom. Vocal stereotypy and singing behavior in baiomyine mice. Journal of Mammalogy, 88(6):1447–1465, 2007.

[34] David Nicholson and Yarden Cohen. vak 0.3. https://doi.org/10.5281/zenodo.4316068, 2020.

[35] Nicolas Stephen Novakowski. The influence of vocalization on the behavior of beaver, castor canadensis kuhl. American Midland Naturalist, pages 198–204, 1969.

[36] Aaron van den Oord, Sander Dieleman, Heiga Zen, Karen Simonyan, Oriol Vinyals, Alex Graves, Nal Kalch-brenner, Andrew Senior, and Koray Kavukcuoglu. Wavenet: A generative model for raw audio. arXiv preprint arXiv:1609.03499, 2016.

[37] Adam Paszke, Sam Gross, Soumith Chintala, Gregory Chanan, Edward Yang, Zachary DeVito, Zeming Lin, Alban Desmaison, Luca Antiga, and Adam Lerer. Automatic differentiation in pytorch. 2017.

[38] Jonathan F Prather, Stephen Nowicki, Rindy C Anderson, Susan Peters, and Richard Mooney. Neural correlates of categorical perception in learned vocal communication. Nature neuroscience, 12(2):221, 2009.

[39] Danilo Jimenez Rezende, Shakir Mohamed, and Daan Wierstra. Stochastic backpropagation and variational inference in deep latent gaussian models. arXiv preprint arXiv:1401.4082, 2014.

[40] Peter J Rousseeuw. Silhouettes: a graphical aid to the interpretation and validation of cluster analysis. Journal of computational and applied mathematics, 20:53–65, 1987.

[41] Monika Sadananda, Markus Wöhr, and Rainer KW Schwarting. Playback of 22-khz and 50-khz ultrasonic vocalizations induces differential c-fos expression in rat brain. Neuroscience letters, 435(1):17–23, 2008.

[42] Tim Sainburg, Brad Theilman, Marvin Thielk, and Timothy Q Gentner. Parallels in the sequential organization of birdsong and human speech. Nature communications, 10, 2019.

[43] Tim Sainburg, Marvin Thielk, and Timothy Q Gentner. Finding, visualizing, and quantifying latent structure across diverse animal vocal repertoires. PLoS computational biology, 16(10):e1008228, 2020.

[44] W John Smith, Sharon L Smith, Elizabeth C Oppenheimer, and Jill G Devilla. Vocalizations of the black-tailed prairie dog, cynomys ludovicianus. Animal behaviour, 25:152–164, 1977.

[45] Kihyuk Sohn, Honglak Lee, and Xinchen Yan. Learning structured output representation using deep conditional generative models. In Advances in neural information processing systems, pages 3483–3491, 2015.

[46] Roland Sossinka and Jörg Böhner. Song types in the zebra finch poephila guttata castanotis. Zeitschrift für Tierpsychologie, 53(2):123–132, 1980.

[47] Daniel Soudry, Suraj Keshri, Patrick Stinson, Min-hwan Oh, Garud Iyengar, and Liam Paninski. Efficient” shotgun” inference of neural connectivity from highly sub-sampled activity data. PLoS computational biology, 11(10):e1004464, 2015.

[48] O Tchernichovski and PP Mitra. Sound analysis pro user manual. CCNY, New York, 2004.

[49] Ofer Tchernichovski, Fernando Nottebohm, Ching Elizabeth Ho, Bijan Pesaran, and Partha Pratim Mitra. A procedure for an automated measurement of song similarity. Animal behaviour, 59(6):1167–1176, 2000.

[50] Maarten Van Segbroeck, Allison T Knoll, Pat Levitt, and Shrikanth Narayanan. Mupet—mouse ultrasonic profile extraction: a signal processing tool for rapid and unsupervised analysis of ultrasonic vocalizations. Neuron, 94(3):465–485, 2017.

[51] J Craig Venter, Mark D Adams, Granger G Sutton, Anthony R Kerlavage, Hamilton O Smith, and Michael Hunkapiller. Shotgun sequencing of the humangenome, 1998.

[52] Alex H Williams, Ben Poole, Niru Maheswaranathan, Ashesh K Dhawale, Tucker Fisher, Christopher D Wilson, David H Brann, Eric Trautmann, Stephen Ryu, Roman Shusterman, et al. Discovering precise temporal patterns in large-scale neural recordings through robust and interpretable time warping. BioRxiv, page 661165, 2019.

[53] Markus Woehr. Ultrasonic vocalizations in shank mouse models for autism spectrum disorders: detailed spectro-graphic analyses and developmental profiles. Neuroscience & Biobehavioral Reviews, 43:199–212, 2014.

